# The DEAD box RNA helicase DDX42 is an intrinsic inhibitor of positive-strand RNA viruses

**DOI:** 10.1101/2020.10.28.359356

**Authors:** Boris Bonaventure, Antoine Rebendenne, Francisco Garcia de Gracia, Joe McKellar, Ségolène Gracias, Emmanuel Labaronne, Marine Tauziet, Ana Luiza Chaves Valadão, Eric Bernard, Laurence Briant, Nathalie Gros, Wassila Djilli, Mary Arnaud-Arnould, Valérie Courgnaud, Hugues Parrinello, Stéphanie Rialle, Emiliano Ricci, Nolwenn Jouvenet, Reiner Schulz, Olivier Moncorgé, Caroline Goujon

**Affiliations:** IRIM, CNRS, Université de Montpellier, France; Institut Pasteur, Virus Sensing and Signaling Unit, Department of Virology, CNRS UMR 3569, Paris, France; LBMC, Université de Lyon, ENS de Lyon, CNRS, INSERM, Lyon, France; CEMIPAI, CNRS, Université de Montpellier, France; IGMM, CNRS, Université de Montpellier, France; Montpellier GenomiX (MGX), Biocampus, CNRS, INSERM, Université de Montpellier, France; Department of Medical & Molecular Genetics, King’s College London, United Kingdom

## Abstract

Genome-wide screens are powerful approaches to unravel new regulators of viral infections. Here, we used a CRISPR/Cas9 screen to reveal new HIV-1 inhibitors. This approach led us to identify the RNA helicase DDX42 as an intrinsic antiviral inhibitor. DDX42 was previously described as a non-processive helicase, able to bind RNA secondary structures such as G-quadruplexes, with no clearly defined function ascribed. Our data show that depletion of endogenous DDX42 significantly increased HIV-1 DNA accumulation and infection in cell lines and primary cells. DDX42 overexpression inhibited HIV-1, whereas a dominant-negative mutant increased infection. Importantly, DDX42 also restricted retrotransposition of LINE-1, infection with other retroviruses and positive-strand RNA viruses, including CHIKV and SARS-CoV-2. However, DDX42 did not inhibit infection with three negative-strand RNA viruses, arguing against a general, unspecific effect on target cells, which was confirmed by RNA-seq analysis. DDX42 was found in the vicinity of viral elements by proximity ligation assays, and cross-linking RNA immunoprecipitation confirmed a specific interaction of DDX42 with RNAs from sensitive viruses. This strongly suggested a direct mode of action of DDX42 on viral ribonucleoprotein complexes. Taken together, our results show for the first time a new and important role of DDX42 in intrinsic antiviral immunity.

## Introduction

The intrinsic and innate immunity are at the frontline against viral invasion and provide a rapid and global defence. The innate immunity relies on viral sensing by Pathogen Recognition Receptors (PRRs) inducing the production of type 1 and 3 interferons (IFNs). Secreted IFNs bind to specific receptors and activate the JAK-STAT signalling cascade, which leads to the expression of hundreds of IFN-stimulated genes (ISGs). The cellular reprogramming induced by ISG expression allows the establishment of an antiviral state that efficiently limits viral replication. Some ISGs are indeed direct antiviral effectors harbouring powerful antiviral activities (1). In addition to the IFN response, antiviral proteins that are constitutively expressed are able to immediately counteract incoming virus replication and are referred to as intrinsic inhibitors; they are part of the so-called intrinsic immunity.

Intensive efforts have been made over the past decades to identify genes able to limit viral replication. Several ISGs were identified a long time ago as major players of innate immunity against viruses, such as the myxovirus resistance protein 1 (MX1) Dynamin Like GTPase, 2′,5′-oligoadenylate synthetases (OASs) and ribonuclease L (RNASeL), or protein kinase R (PKR) (2–4). More recently, gain-of-function and loss-of-function screens have identified new IFN-induced antiviral effectors (5–7). A growing list of cellular proteins with various functions has hence been identified as capable of limiting different steps of virus life cycles (8–10). Viruses have often evolved to counteract the action of these so-called restriction factors. However, type 1 IFNs (e.g. IFN-alpha and -beta) induce, through the expression of ISGs, an antiviral state particularly efficient at inhibiting HIV-1 when cells are pre-exposed to IFN (9). The dynamin-like GTPase MX2, and, recently, the restriction factor TRIM5α, have both been shown to participate in the IFN-induced inhibition of HIV-1 (6, 11–13). While numerous antiviral ISGs have been identified, less is probably known about the extent of the intrinsic, antiviral inhibitor repertoire. The recent identification of TRIM7 as an intrinsic inhibitor of enteroviruses illustrates the fact that important antiviral inhibitors most certainly remain to be revealed (14).

With the hypothesis that additional HIV-1 inhibitors remained to be identified, we took advantage of the hostile environment induced by IFN to develop a whole-genome, CRISPR/Cas9 screen strategy in order to reveal intrinsic and innate inhibitors. This strategy led us to identify DDX42 as a new intrinsic inhibitor of HIV-1, acting independently of the IFN system. We reveal that endogenous DDX42 is antiviral in various cell types, including primary targets of HIV-1, and impairs the accumulation of viral DNA. Moreover, our data show a broad activity against lentiviruses and the retrovirus Murine Leukemia Virus (MLV). Reminiscent of other HIV-1 inhibitors such as APOBEC3G, DDX42 also blocks LINE-1 spread by interacting with their RNAs. Interestingly, while three different negative strand RNA viruses were found insensitive to DDX42, several positive strand RNA viruses, including the flavivirus Zika (ZIKV), the alphavirus Chikungunya (CHIKV) and the severe acute respiratory syndrome coronavirus 2 (SARS-CoV-2), were inhibited by this non-processive RNA helicase. Finally, cross-linking RNA immunoprecipitation assays showed that DDX42 specifically binds to viral RNAs from sensitive viruses, strongly suggesting a direct mode of action. Overall, our study sheds light on a new intrinsic, antiviral function of a so far poorly studied DEAD-box RNA helicase, and provide new insights on a broad-spectrum antiviral inhibitor.

## Results

The Genome-Scale CRISPR Knock-Out (GeCKO) sgRNA library (15–17) was used to generate knock-out (KO) populations in the glioblastoma T98G cell line. This cell line is both highly permissive to HIV-1 infection and potently able to suppress infection following type 1 IFN treatment (Fig. S1A). The screen strategy is depicted in Figure 1A. Cas9-expressing T98G cells were independently transduced with lentiviral vectors (LVs) coding the two-halves of the GeCKO library, at a low multiplicity of infection (MOI). Next-generation sequencing showed more than 94% sgRNA coverage for each sub-library (not shown). Cells were pre-treated with type 1 IFN (IFN-alpha) and incubated with VSV-G-pseudotyped, HIV-1 based LVs coding for an antibiotic resistance cassette. The cells which were successfully infected despite the IFN treatment were selected by survival in the presence of antibiotics. In order to enrich the populations with mutants of interest and to limit the presence of false-positives, two additional rounds of IFN treatment, infection and selection (with different antibiotics) were performed. As expected, the cells enriched after each round became less refractory to HIV-1 infection following IFN treatment (Fig. S1B).

**Figure 1.**
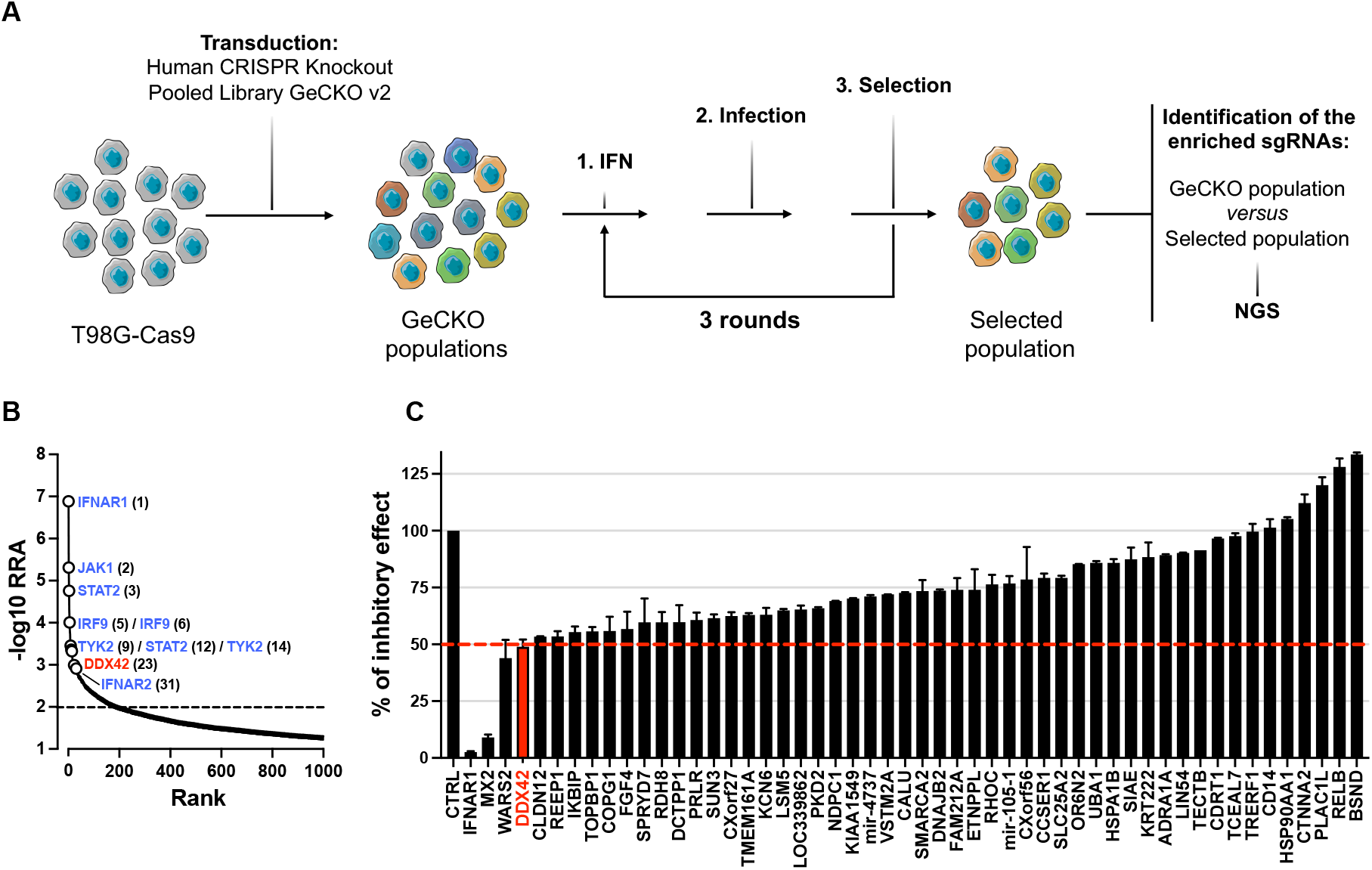
A whole-genome CRISPR/Cas9 screen to identify new HIV-1 inhibitors. **A.** Screen strategy. GeCKO cell populations (obtained by transduction of T98G/Cas9 cells with GeCKO v2 LV library) were IFN-treated, challenged with HIV-1 LVs coding for an antibiotic resistance gene and selected. Three rounds of IFN treatment, infection and selection were performed. Genomic DNAs of initial GeCKO and 3-time selected populations were extracted, the sgRNA-coding sequences amplified and sequenced. **B.** Candidate gene identification. MAGeCK computational statistical tool (18) was used to establish a Robust Rank Aggregation (RRA) score for each gene based on sgRNA enrichment and number of sgRNAs per gene. Genes belonging to the type 1 IFN response pathway (in blue) and DDX42 (in red) are shown (respective ranks into brackets) for 2 independent screens (the results of which were merged in the analysis). **C.** Candidate validation. T98G/Cas9/CD4/CXCR4/Firefly KO populations were generated for the 25 top hits of each screen. The control (CTRL) condition represents the mean of 4 negative CTRL populations, obtained with 4 non-targeting sgRNAs; *IFNAR1* and *MX2* KO populations were used as positive controls. KO cell populations were treated with IFN and infected with HIV-1 Renilla and luciferase signals were measured 30 h later (Renilla signals were normalized to Firefly). IFN inhibition (i.e. ratio of untreated / IFN-treated conditions) was calculated and set at 100% inhibition for CTRL. A representative experiment is shown (mean and standard deviation from technical duplicates).

The differential sgRNA abundance between the initial GeCKO populations and selected populations was analysed by next-generation sequencing (NGS) and the MAGeCK algorithm was used to rank the candidate genes (Fig. 1B). An enrichment was observed for 200 genes (RRA score > 0,01), with the best hits being *IFNAR1*, *JAK1* and *STAT2* (Fig. 1B). The crucial mediators of type 1 IFN signalling cascade were among the top hits in both screens (with the notable exception of *STAT1*), validating our approach and confirming the identification of relevant genes. Interestingly, most of the other candidates displayed unknown functions, or functions that were *a priori* unrelated to innate immunity. Of note, very little overlap was observed between the two independent screens, performed with two sub-libraries. However, such a poor overlap between biological replicates has been observed before and does not preclude obtaining valid data (19). The top 25 candidate genes from each screen were selected for further validation. T98G/Cas9 cells expressing HIV-1 CD4 and CXCR4 receptors, as well as Firefly luciferase as an internal control (T98G/Cas9/CD4/CXCR4/Firefly cells), were transduced with sgRNA-expressing LVs to generate individual KO populations, using the identified sgRNA sequences. Four irrelevant, non-targeting sgRNAs, as well as sgRNAs targeting *IFNAR1* and *MX2*, were used to generate negative and positive control populations, respectively. The KO cell populations were pre-treated with IFN and infected with an HIV-1 reporter virus expressing Renilla luciferase and bearing HIV-1 envelope (20) (hereafter called HIV-1 Renilla). Infection efficiency was analysed 30 h later (Fig. 1C). As expected (11, 21, 22), *IFNAR1* and *MX2* KO fully and partially rescued HIV-1 infection from the protective effect of IFN, respectively. The KO of two candidate genes, namely *WARS2* and *DDX42*, allowed a partial rescue of HIV-1 infection from the IFN-induced inhibition, suggesting a potential role of these candidate genes in HIV-1 inhibition.

DDX42 is a member of the DEAD box family of RNA helicases, with RNA chaperone activities (23) and, as such, retained our attention. Indeed, DEAD box helicases are well-known to regulate HIV-1 life cycle (24). However, to our knowledge, the impact of DDX42 on HIV-1 replication had never been studied. In order to validate the effect of *DDX42* KO on HIV-1 infection in another model cell line, two additional sgRNAs were designed (sgRNA-2 and −3) and used in parallel to the one identified in the GeCKO screen (sgDDX42-1) (Fig. 2A). U87-MG/CD4/CXCR4 cells were used here, as we previously extensively characterized the IFN phenotype in these cells (11). Control and *DDX42* KO cell populations were generated and pre-treated or not with IFN prior to infection with HIV-1 Renilla. Of note, CRISPR/Cas9 KO of *DDX42* induced only a partial decrease of DDX42 protein levels (Fig. 2A) and cell populations tended to derive (not shown), suggesting a potential role for DDX42 in cell proliferation or long-term survival. We observed however that DDX42 partial depletion with all 3 sgRNAs improved HIV-1 infection, confirming that endogenous DDX42 had a negative impact on HIV-1 replication. Interestingly, the increase in infection efficiency induced by *DDX42* KO was observed independently of the IFN treatment. DDX42 is not an ISG, as shown in several cell lines (e.g. U87-MG, T98G, HEK293T) and in primary T cells and monocyte-derived macrophages (Fig. S2A and GSE46599 (11)). The fact that the IFN-induced state is at least partially saturable (Fig. S1A) explains why an intrinsic inhibitor of HIV-1, which is not regulated by IFN, could be identified by our approach. Indeed, removing one barrier to infection presumably rendered the cells more permissive and, in this context, IFN had less of an impact on viral replication.

**Figure 2.**
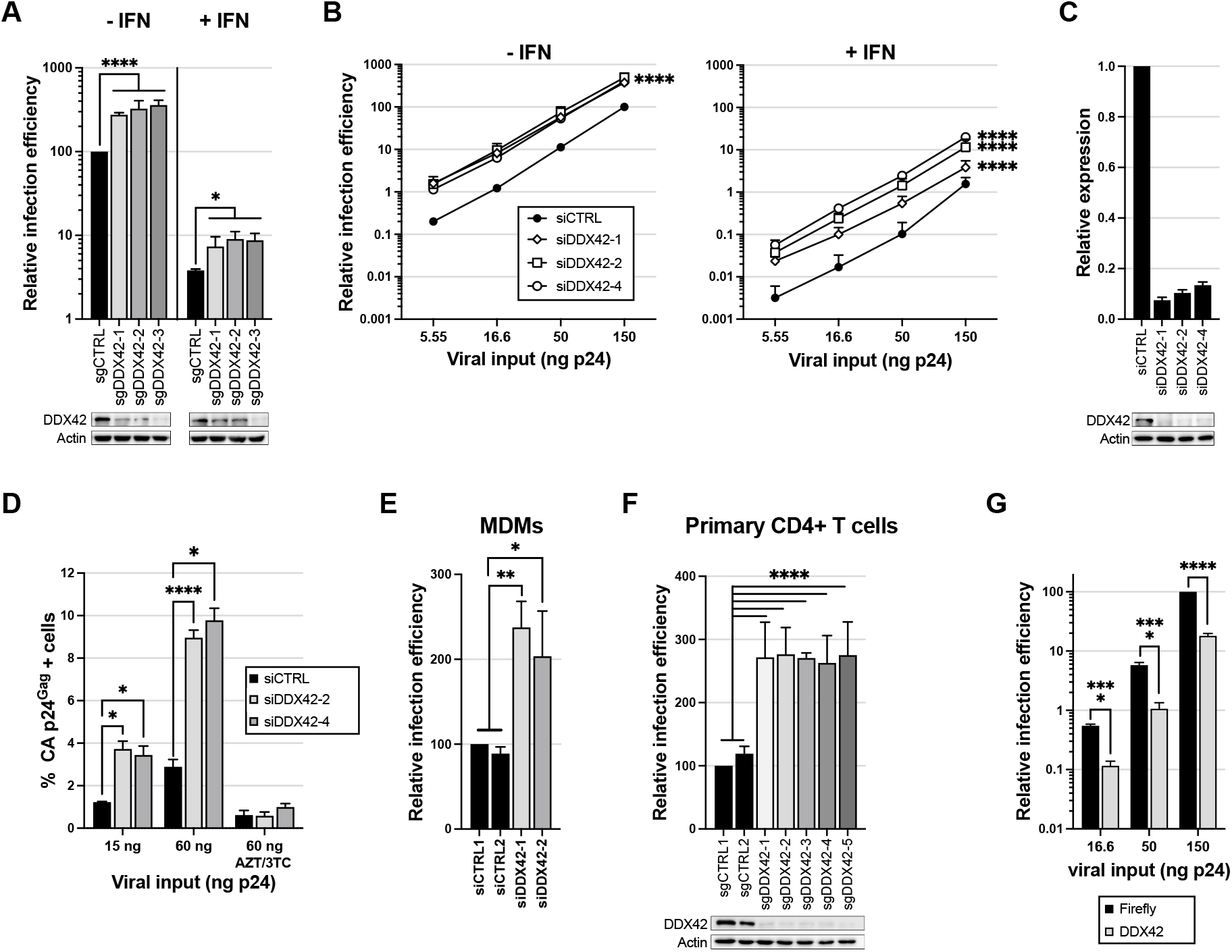
DDX42 is an intrinsic inhibitor of HIV-1. **A.** Top: *DDX42* KO and CTRL KO U87-MG/CD4/CXCR4/Cas9/Firefly cells were generated using 3 sgRNAs and 4 non-targeting sgRNAs, respectively (for CTRL, the average of data obtained with 4 cell populations is shown). Cells were treated or not with IFN 24 h prior to infection with HIV-1 Renilla. Relative luminescence results for IFN-treated and -untreated conditions are shown. Two-way ANOVA on log-transformed data with Sidak’s test. Bottom: Immunoblot analysis of DDX42 levels is shown for 1 CTRL and DDX42-depleted populations; Actin served as a loading control. **B.** siRNA-transfected U87-MG/CD4/CXCR4 cells were treated or not with IFN for 24 h prior to infection with HIV-1 Renilla. Relative luminescence results for IFN-treated and -untreated conditions are shown. Multiple linear regression analysis. **C.** DDX42 silencing efficiency measured by RT-qPCR (top) and immunoblot (bottom) in parallel samples from B. **D.** DDX42-depleted cells were infected with HIV-1, and infection efficiency was measured by CA p24^Gag^ intracellular staining and flow cytometry analysis. When indicated, cells were treated with azidothymidine (AZT) and lamivudine (3TC). Two-way ANOVA on log-transformed data with Dunnett’s test. **E.** siRNA-transfected MDMs were infected with a CCR5-tropic version of HIV-1 Renilla. Relative luminescence results from independent experiments performed with cells from three donors are shown. Two-way ANOVA on log-transformed data with Dunnett’s test. **F.** Primary CD4+ T cells were electroporated with Cas9-sgRNA RNPs using 2 non-targeting sgRNAs (sgCTRL1 and 2) and 5 sgRNAs targeting DDX42. Top: Cells were then infected with HIV-1 Renilla and relative infection efficiencies obtained with cells from three donors are shown. Two-way ANOVA on log-transformed data with Dunnett’s test. Bottom: DDX42 protein levels were determined by immunoblot, Actin served as a loading control. A representative immunoblot is shown. **G.** Firefly- or DDX42-expresssing U87-MG/CD4/CXCR4 cells were infected with HIV-1 Renilla. Relative infection efficiencies are shown. Multiple linear regression analysis. **A-G** Data represent the mean ± S.E.M of three independent experiments.

In order to confirm DDX42’s effect on HIV-1 infection with an independent approach, we used three different siRNAs to knockdown DDX42 expression. DDX42 depletion increased HIV-1 Renilla infection efficiency by 3 to 8-fold in U87-MG/CD4/CXCR4 cells, irrespectively of the presence of IFN (Fig. 2B). Of note, DDX42 depletion was highly efficient (~90% efficiency both at the mRNA and protein levels, Fig. 2C) but had no effect on cell viability (Fig. S2B). Importantly, wild-type HIV-1 infection was also significantly impacted by DDX42 silencing, as measured by Capsid (CA p24^Gag^) intracellular staining 30 h post-infection in U87-MG/CD4/CXCR4 cells (Fig. 2D). We then investigated whether DDX42 had an impact in HIV-1 primary target cells. In monocyte-derived macrophages (MDMs), we observed that HIV-1 infection was increased by about 2-fold following DDX42 silencing (Fig. 2E), whereas DDX42 mRNA abundance was decreased by only 40% (Fig. S2C). Electroporation of pre-assembled Cas9-sgRNA ribonucleoprotein complexes (RNPs) was used to efficiently deplete DDX42 in primary CD4+ T cells (Fig. 2F). DDX42 depletion increased HIV-1 infection by 2- to 3-fold, showing a physiological role of DDX42 as an intrinsic inhibitor of HIV-1 in primary CD4+ T cells. Next, we analysed the consequences of DDX42 overexpression on HIV-1 infection. An irrelevant control (Firefly) or DDX42 were ectopically expressed in U87-MG/CD4/CXCR4 and the cells were challenged with HIV-1 (Fig. 2G). DDX42 overexpression induced a substantial inhibition of HIV-1 infection (~5-fold decrease in infection efficiency in comparison to the control). Interestingly, the expression of K303E DDX42 mutant, which is unable to hydrolyse ATP and may supposedly act as a dominant negative (25, 26), increased HIV-1 infection by 3-fold (Fig. S2D), reminiscent of the impact of DDX42 depletion. Altogether, these data showed that endogenous DDX42 is able to intrinsically inhibit HIV-1 infection.

In order to determine which step of HIV-1 life cycle was affected by DDX42, we quantified HIV-1 DNA accumulation over time in DDX42-silenced and control cells (Fig. 3A; silencing efficiency is shown in Fig. 3B). DDX42 depletion increased accumulation of early and late reverse transcript products (by 2.5- to 8-fold), as well as proviral DNA and 2-long terminal repeat (2-LTR) circles at 48h post-infection (by 2.5- to 4.5-fold). Importantly, DDX42 silencing did not impact HIV-1 entry (Fig. S3A). These data suggested that endogenous DDX42 could inhibit reverse transcription and/or impact genome stability, leading to a decrease in viral DNA accumulation. We hypothesized that if that was the case, DDX42 should be found in close proximity to HIV-1 reverse transcription complexes during infection. In agreement with this, proximity ligation assay (PLA) performed on HIV-1 infected MDMs showed that DDX42 was indeed in close vicinity of Capsid (Fig. 3C).

**Figure 3.**
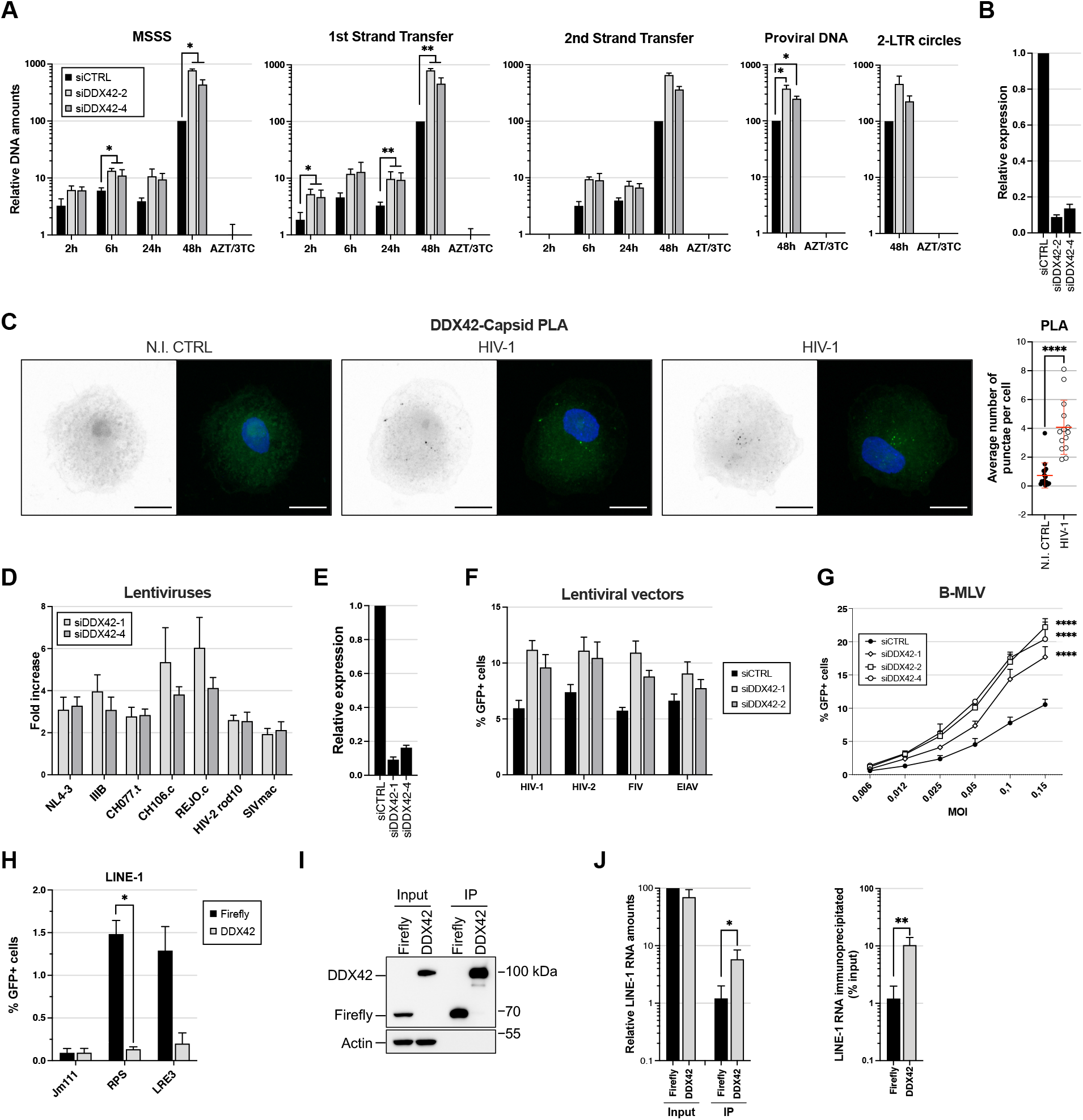
Characterization of DDX42 inhibitory activity against retroviruses and retroelements. **A.** siRNA-transfected U87-MG/CD4/CXCR4 cells were infected with HIV-1 and relative amounts of Minus Strand Strong Stop (MSSS), 1^st^ and 2^nd^ Strand Transfer DNAs, and nuclear forms of HIV-1 DNA (proviral DNA, and 2-LTR circles) were quantified by qPCR. DNAs from cells infected for 48 h in the presence of AZT and 3TC were used as a control. Mixed-effects analysis on log-transformed data with Dunnett’s test. **B.** Silencing efficiency in parallel samples from A. **C.** PLAs were performed in MDMs infected with HIV-1 or not (N.I. CTRL), using anti-Capsid and anti-DDX42 antibodies (nuclei stained with Hoechst). Images were acquired using a LSM880 Airyscan microscope. Left: representative images, scale-bar: 10 μm. Right: Average punctae quantified per cell in 3 independent assays done on MDMs from different donors with mean ± SD (n>65 cells per condition). Mann-Whitney test. **D.** siRNA-transfected TZM-bl cells were infected and β-galactosidase signals measured 24 h later. The ratio of the signal in DDX42-depleted versus CTRL cells is shown. **E.** Silencing efficiency in parallel samples from D. **F.** siRNA-transfected U87-MG/CD4/CXCR4 cells were infected with HIV-1- HIV-2- FIV- EIAV-based, GFP-coding LVs and infection efficiency was scored 24 h later by measuring the percentage of GFP expressing cells by flow cytometry. **G.** siRNA-transfected U87-MG/CD4/CXCR4 cells were infected with GFP-coding B-MLV and infection efficiency measured 24h later by flow cytometry. Simple linear regression analysis. **H.** HEK293T were co-transfected with GFP-coding LINE-1 plasmids (RPS-GFP or LRE3-GFP) or with an inactive LINE-1 plasmid (JM111) together with either a Firefly- or DDX42-coding plasmid. GFP expression was measured by flow cytometry 7 days later. Two-way ANOVA on log-transformed data with Sidak’s test. **I.** HEK293T were co-transfected with pRPS-GFP and a Flag-Firefly- (negative control) or Flag-DDX42-coding plasmid, followed by Flag immunoprecipitation and immunoblot analysis. A representative immunoblot is shown. **J.** Left, RNA extraction and LINE-1 RT-qPCR on parallel samples from H. Two-way ANOVA on log-transformed data with Sidak’s test. Right, Percentage of immunoprecipitated RNA from I. T-test on log-transformed data. **A-H.** Data represent the mean ± S.E.M of 3 independent experiments. **J.** Data represent the mean ± S.E.M of 5 independent experiments.

We next examined the ability of DDX42 to inhibit infection by various primate lentiviruses, including lab-adapted strains of HIV-1 (NL4-3, IIIB), transmitted/founder strains (27) (CH077.t, CH106.c, REJO.c), HIV-2 and simian immunodeficiency virus from rhesus macaque (SIVMAC). DDX42 was depleted or not in TZM-bl reporter cells prior to infection with VSV-G-pseudotyped lentiviruses, and infection efficiency was monitored 24h later (Fig. 3D; silencing efficiency is shown in Fig. 3E). DDX42 depletion increased infection levels similarly with all the tested strains of HIV-1 (i.e. 3- to 5-fold). HIV-2rod10 and SIVMAC infection efficiencies were also slightly improved in the absence of DDX42 (~2-fold). The analysis was then extended to two non-primate lentiviruses, equine infectious anemia virus (EIAV) and feline immunodeficiency virus (FIV), using GFP-coding LVs in U87-MG cells (Fig. 3F). DDX42 depletion appeared to increase HIV-1, HIV-2 and FIV LV infection to the same extent (~2-fold), whereas EIAV infection was less impacted. Of note, DDX42 antiviral activity appeared less potent on HIV-1 LVs compared to full-length HIV-1, which might suggest that genome length or cis-acting elements could play a role in DDX42 inhibition. We also observed that DDX42 depletion led to a significant increase in infection with GFP-coding, MLV-derived vectors (Fig. 3G). These results strongly support a general antiviral activity of DDX42 against retroviruses.

DDX42 can be found in the cytoplasm but is predominantly located in the nucleus in various cell types, including monocyte-derived macrophages (28, 29) (Fig. S3B). Considering that DDX42 showed a broad activity against retroviruses and seemed to act at the level of reverse transcription, we sought to investigate whether DDX42 could inhibit retrotransposons. Long interspersed nuclear elements (LINE)-1 are non-LTR retrotransposons, which have been found to be active in the germ line and some somatic cells (30). Interestingly, DDX42 was identified among the suppressors of LINE-1 retrotransposition through a genome-wide screen in K562 cells, although not further characterized (31). To confirm that DDX42 could inhibit LINE-1 retrotransposition, HEK293T cells were co-transfected with GFP-expressing LINE-1 plasmids (RPS or LRE3) or an inactive LINE-1 (JM111) together with a DDX42- or a control (Firefly)-expressing plasmid (32). LINE-1 retrotransposition was quantified by flow cytometry 7 days later (Fig. 3H). As the GFP cassette is cloned in antisense and disrupted by an intron, GFP is only expressed after LINE-1 transcription, splicing, Orf2p-mediated reverse transcription, and integration (32). Successful retrotransposition events were observed in >1.25% of control cells, but in only <0.25% of DDX42-expressing cells (i.e. a percentage similar to what observed with the non-active LINE-1), showing that DDX42 ectopic expression significantly suppressed LINE-1 retrotransposition. Next, we investigated whether DDX42 could physically interact with LINE-1 RNAs by performing cross-linking RNA immunoprecipitation. Cells were co-transfected with GFP-expressing LINE-1 RPS plasmid and either flag-tagged-Firefly or -DDX42. Four days later, the cells were treated with formaldehyde, lysed and the flagged proteins immunoprecipitated. The immunoprecipitation eluates were then divided in two; the immunoprecipitated proteins were analysed by immunoblot (Fig. 3I) and their associated RNAs were extracted and analysed by RT-qPCR using LINE-1 specific primers (Fig. 3J). A significant enrichment of LINE-1 RNAs was observed with DDX42 immunoprecipitation as compared to the Firefly negative control, showing that DDX42 could interact with LINE-1 RNAs.

Finally, we sought to determine whether DDX42’s inhibitory activity was specific towards retroviruses and retroelements, or could be extended to other viruses, as observed for many other anti-HIV-1 proteins, such as ZAP or BST-2/Tetherin (1, 8). To this aim, we tested the impact of DDX42 depletion on nine RNA viruses from six different families: the orthomyxovirus influenza A virus (IAV), the rhabdovirus vesicular stomatitis virus (VSV), the paramyxovirus measles virus (MeV), the flaviviruses ZIKV, Dengue virus serotype 2 (DENV-2) and yellow fever virus (YFV), the alphavirus CHIKV, the coronavirus SARS-CoV-2, which is responsible for the current coronavirus disease (COVID)-19 pandemic, and the seasonal coronavirus HCoV-229E (Fig. 4 and S4). Strikingly, DDX42 depletion had no significant effect on IAV, VSV and MeV replication in U87-MG and Huh-7 cells, respectively (Fig. 4A-C; silencing efficiency is shown in Fig. S4A), thereby strongly suggesting that manipulating DDX42 expression did not have a broad and unspecific impact on target cells. Interestingly, depletion of endogenous DDX42 had a modest but significant, positive impact on ZIKV in U87-MG cells (Fig. 4D), whereas the impact in Huh-7 cells was minor (Fig. S4B and A), probably due to the fact that the silencing was less efficient in the latter or, alternatively, that a wild-type version of ZIKV and a different readout was used in this case. Similarly, DDX42 silencing had little effect on 2 other flaviviruses (DENV-2 and YFV) in Huh-7 cells, as measured by the number of cells positive for the viral protein E (Fig. S4C-D). By contrast, DDX42 depletion had a profound effect on CHIKV (Fig. 4E), SARS-CoV-2 (Fig. 4F-G) and HCoV-229E (Fig 4SA and E) replication (up to 1 log-increase in infection efficiency with CHIKV and HCoV-229, and 3 log-increase with SARS-CoV-2; silencing efficiencies in the different cell lines used are shown in Fig. S4A). Plaque assays confirmed a strong impact of DDX42 depletion on infectious SARS-CoV-2 production (Fig. 4G). In agreement with this, DDX42 was recently identified as a potential inhibitor of SARS-CoV-2 replication in a whole-genome CRISPR screen in simian cells (33). Next, we used PLA to determine whether DDX42 was in the vicinity of SARS-CoV-2 components in infected cells. To this aim, PLA was performed with either an anti-double strand (ds)RNA or anti-Nucleoprotein

**Figure 4.**
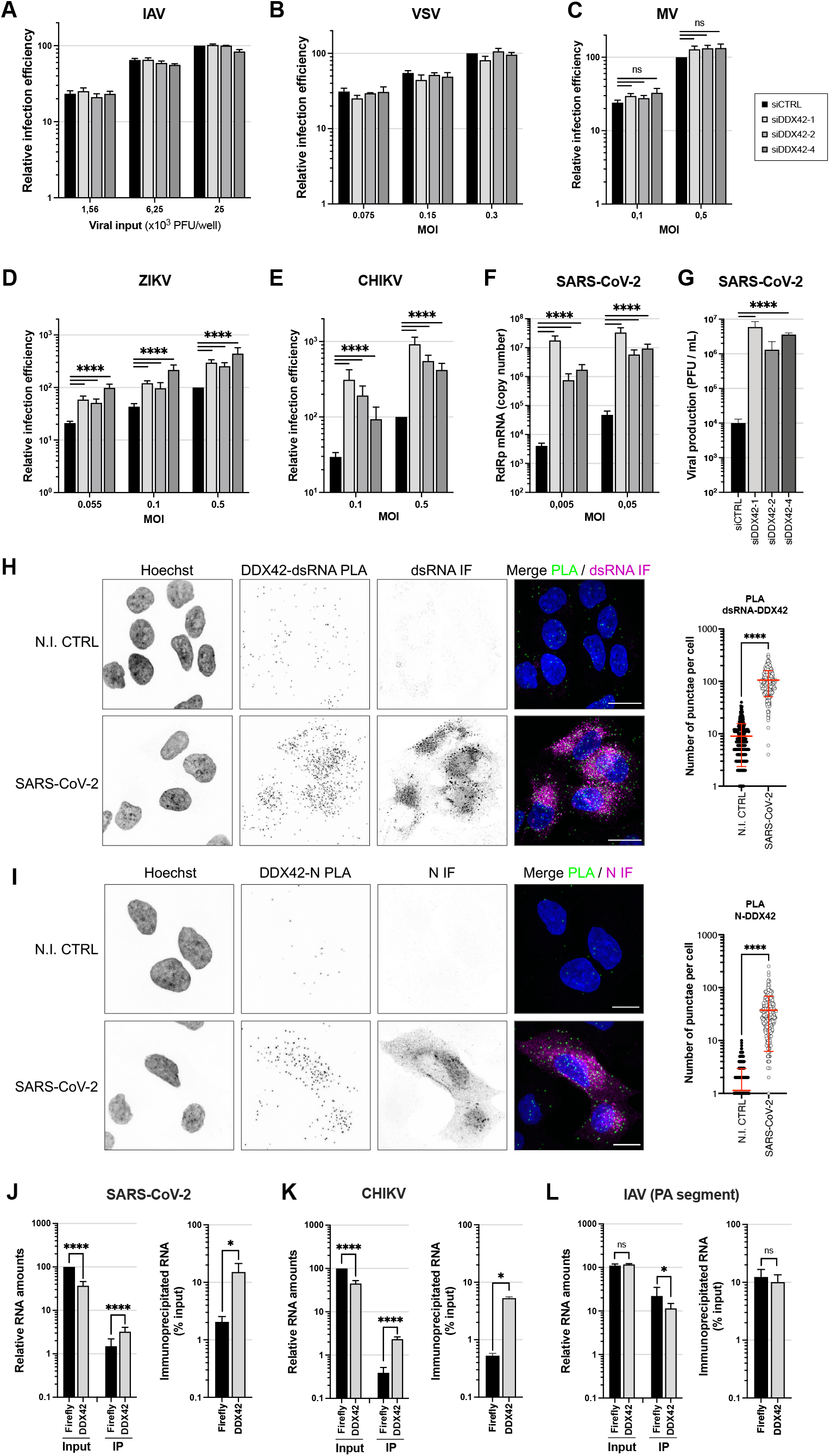
DDX42 exerts a broad antiviral activity on positive strand RNA viruses and interacts with viral RNAs from targeted viruses. **A.** Relative IAV infection efficiency in siRNA-transfected U87-MG cells (Nanoluciferase activity 16 h post infection). **B.** Relative VSV infection efficiency in siRNA-transfected U87-MG cells (Firefly activity 24 h post infection). **C.** Relative MeV infection efficiency in siRNA-transfected Huh-7 cells (GFP+ cells scored 24h post infection). Multiple linear regression analysis. **D.** Relative ZIKV infection efficiency in siRNA-transfected U87-MG cells (Nanoluciferase activity 24 h post infection). Multiple linear regression analysis. **E.** Relative CHIKV infection efficiency in siRNA-transfected U87-MG cells (Nanoluciferase activity 24 h post infection). Multiple linear regression analysis. **F.** Relative SARS-CoV-2 infection in siRNA-transfected A549-ACE2 cells (RdRp RT-qPCR 48 h post-infection). Mixed-effects analysis on log-transformed data with Dunnett’s test. **G.** Viral production in cell supernatants from F (48 h post-infection, MOI 0,05) measured by plaque assays. Two-way ANOVA on log-transformed data with Dunnett’s test. **H.** A549-ACE2 cells were infected or not with SARS-CoV-2 for 24 h prior to PLA using mouse anti-dsRNA (J2) and rabbit anti-DDX42 antibodies, followed by additional immunofluorescence (IF) staining with anti-mouse Alexa Fluor 546 antibody (PLA in green, IF in magenta). Representative Z-stack projection images are shown; scale bar: 15 μm. Average punctae were quantified in 3 independent PLA assays with mean ± SD (n>75 cells per condition). Mann-Whitney test. **I.** Identical to H but using an anti-N antibody instead of anti-dsRNA antibody. **J.** Left, Quantification of SARS-CoV-2 RNA by RT-qPCR in RNA from total U87-MG-ACE2 cell lysates (input) and in Flag-Firefly (negative control) and Flag-DDX42 immunoprecipitation (IP). Multiple linear regression analysis on log-transformed data. Right, Percentage of immunoprecipitated RNA. Paired t-test on log-transformed data. **K.** Identical to J following CHIKV infection of U87-MG cells. **L.** Identical to J following IAV infection of U87-MG cells. Data represent the mean ± S.E.M. of three (**A-E, J-L**), four (**F**) or two (**G**) independent experiments.

(N) antibody, together with an anti-DDX42 antibody, followed by immunofluorescence staining to identify the infected cells (Fig. 4H-I). In the latter, there was a significantly higher number of dsRNA-DDX42 and N-DDX42 PLA punctae than in control cells. This proximity suggested a potential interaction between DDX42 and SARS-CoV-2 viral components. To test whether DDX42 RNA helicase could indeed interact with viral RNAs, cross-linking RNA immunoprecipitation experiments were conducted following viral infection of U87-MG cells expressing either Flag-DDX42 or negative control Flag-Firefly (in addition to ACE2 for SARS-CoV-2 infection). A significant enrichment of the viral RNAs recovered with Flag-DDX42 was observed in comparison to the negative control. Importantly, such an enrichment was observed for the sensitive viruses SARS-CoV-2 and CHIKV (Fig. 4J-K), but not for IAV, which was insensitive to DDX42 antiviral activity (Fig. 4L and 4A).

As mentioned above, DDX42’s lack of effect on negative strand RNA viruses argued against a global, indirect impact on the target cells. However, to confirm this, we performed RNA-seq analysis on siRNA-treated U87-MG and A549-ACE2 cells. The results showed that DDX42 depletion didn’t have a substantial impact on global cellular RNA expression (Supplementary File 1 and Fig. S5). Of note, only 63 genes were commonly found differentially expressed upon DDX42 depletion with the 3 different siRNAs in U87-MG cells, and only 23 differentially expressed genes (DEGs) were identified in common in U87-MG and A549-ACE2 cells. Importantly, no known restriction factors were identified among these DEGs (Supplementary File 1). Taken together, these data strongly suggested that the DEAD-box RNA helicase DDX42 directly impacted viral replication, by interacting with viral RNAs.

## Discussion

Here, we identified for the first time the RNA helicase DDX42 as an intrinsic inhibitor of HIV-1, capable of limiting the efficiency of viral DNA accumulation. Moreover, our study revealed broad activity of endogenous DDX42 against retroviruses and retroelements, which was observed in various cell lines and primary cells, including primary CD4+ T cells. Strikingly, we observed that DDX42 was able to inhibit viruses from other families, which possess different replication strategies, including SARS-CoV-2 and CHIKV. However, DDX42 did not have an impact on all the viruses we tested, as three different negative-strand RNA viruses were found insensitive to DDX42 antiviral activity. This is reminiscent of other broad-spectrum antiviral inhibitors such as MX1, which possesses some specificity despite being able to inhibit viruses from various families (34). Further work is now warranted to explore in depth the breadth of DDX42 antiviral activity and determine whether it is truly specific of positive strand RNA viruses.

Noteworthy, our PLA assays showed a close proximity between DDX42 and HIV-1 Capsid, which is a viral protein recently shown to remain associated with reverse transcription complexes until proviral DNA integration in the nucleus (35–37). We also observed a close proximity between DDX42 and SARS-CoV-2 N or dsRNA, which could be suggestive of a potential interaction of DDX42 with viral components. In line with this, LINE-1 RNAs, as well as SARS-CoV-2 and CHIKV RNAs (but not IAV RNAs), were specifically pulled-down when DDX42 was immunoprecipitated. Taken together, these observations strongly suggest a direct mode of action of DDX42, which could interact with target viral RNAs. Interestingly, DDX42 is known to be a non-processive helicase, which also possesses RNA annealing activities and the ability to displace RNA-binding proteins from single-stranded RNAs (23). Moreover, DDX42 binds to G-quadruplexes (38), which are secondary structures found both in cellular and viral nucleic acids and involved in various processes, such as transcription, translation and replication (39, 40). The functional relevance of this ability to bind G-quadruplexes is currently unknown. Interestingly, DDX42 was initially identified as a transient interactant of the spliceosome component SF3b, and had been proposed to play a role in RNP remodeling in this context, but this was not experimentally tested (41). Whether DDX42 actually plays a role in the splicing machinery formation, or in any other cellular function(s), remains therefore to be assessed. A better knowledge of DDX42 cellular functions may help understanding its mechanism of action against viruses. Of note, the current known activities for DDX42 could be consistent with a potential antiviral role through binding of viral secondary RNA structures and/or through viral RNP remodeling (23, 42).

DEAD box RNA helicases are well-known to participate in innate immunity, with some of the best examples being RIG-I-like receptors, which play an important role in viral nucleic acid sensing (43). Additionally, some RNA helicases are involved in regulating (either positively or negatively) innate immune signaling, while also being involved in various cellular functions. For instance, DDX46 uses both its helicase and ATPase activities to catalyze conformational changes and complex formation in the context of pre-spliceosome assembly (41, 44). Besides this role, DDX46 was shown to negatively regulate innate immune activation by recruiting an m6A eraser onto mRNAs coding key IFN signaling adaptors, such as MAVS, causing their nuclear retention and decreasing IFN production (45). Other RNA helicases are well-known inhibitors of viruses and/or retroelements, like MOV10 for instance. These RNA helicases can generally be IFN-regulated (24). In contrast, there is very few known examples of RNA helicases with antiviral effector activities, which would totally be IFN-independent. To our knowledge, so far, in addition to DDX42, only DDX17 has been shown to possess intrinsic antiviral functions. DDX17 is active against negative strand RNA bunyaviruses, through binding of viral RNA secondary structures, and this activity seemed conserved in Drosophila and humans (43). DDX17 and DDX42 are likely to ensure complementary functions by targeting different viruses, through distinct RNA-binding specificities. Nevertheless, DDX42’s broad spectrum of action seems so far a unique feature among RNA helicases and further work is now needed to understand the viral determinants for DDX42 restriction activity. While G-quadruplexes could be preferential binding sites for DDX42 (23), viral genomes harbor numerous highly structured elements that could also be involved.

In conclusion, this work shows for the first time an antiviral role for DDX42 and highlights the importance of understanding the mechanism of action of this RNA helicase and how it may participate in the control of positive-strand RNA virus replication. Such an understanding might contribute to the development of future antiviral interventional strategies.

## Methods

### Plasmids

The pLentiCas9-Blast, pLentiGuide-Puro vectors and the GeCKO sub-library A and B plasmids were a gift from Prof. F. Zhang (Addgene #52962, #52963, and #1000000048, respectively (17)). LVs coding for sgRNAs targeting the candidate genes and control genes were obtained by cloning annealed oligonucleotides in BsmBI-digested pLentiGuide-Puro, as described (Addgene). Control sgRNAs and sgRNAs targeting the candidate genes, *MX2* and *IFNAR1*, were designed with the Optimized CRISPR Design tool (not available anymore), or with Chopchop (chopchop.cbu.uib.no). The sgRNA coding sequences used were as follow: MX2 5’-CCGCCATTCGGCACAGTGCC-3’, IFNAR1 5’-GACCCTAGTGCTCGTCGCCG-3’, sgCTRL-1 5’-AGCACGTAATGTCCGTGGAT-3’, sgCTRL-2 5’-CAATCGGCGACGTTTTAAAT-3’, sgCTRL-3 5’-TTAATTTGGGTGGGCCCTGC-3’, sgCTRL-4 5’-TTGGATATTAATTAGACATG-3’, sgDDX42-1 5’-TCCTGAACCACACCAGCAGT-3’, sgDDX42-2 5’-GGTGGTCCTGGCACTAAGCG-3’, sgDDX42-3 5’-AGGCACTGTGGGACTGCTGT-3’. All the other sgRNA sequences are available upon request. In order to produce the HIV-1 based LVs used to perform the different steps of the screen (pRRL.sin.cPPT.CMV/NeomycinR.WPRE, pRRL.sin.cPPT.CMV/HygromycinR.WPRE and pRRL.sin.cPPT.CMV/ZeocinR.WPRE), neomycin, hygromycin and zeocin resistance genes (i.e. the genes coding for Neomycin phosphotransferase II, Hygromycin B phosphotransferase, and *Sh ble*) were amplified by PCR from pcDNA3.1 (ThermoFisher Scientific), pAHM (11), and pcDNA3.1/Zeo (ThermoFisher Scientific), respectively, and cloned by replacement of GFP in pRRL.sin.cPPT.CMV/GFP.WPRE (46) using BamHI and SalI restriction sites. The pRRL.sin.cPPT.SFFV/E2-crimson-IRES-PuromycinR.WPRE has been described (47). Human DDX42 cDNA was amplified by RT-PCR using the SuperScript III™ (Invitrogen) from mRNAs of MDMs using primers DDX42-forward 5’-AATTAATTTAGGATCCATGAACTGGAATAAAGGTGGTCCTG and DDX42-reverse 5’-AATTAATTTACTCGAGCTAACTGTCCCATCGACTTTTCTTGCG, and cloned by replacement of E2-crimson in BamHI-XhoI-digested pRRL.sin.cPPT.SFFV/E2crimson-IRES-PuromycinR.WPRE, in order to obtain pRRL.sin.cPPT.SFFV/DDX42-IRES-PuromycinR.WPRE.

The pRRL.sin.cPPT.SFFV/CD4-IRES-CXCR4.WPRE was obtained by replacement of E2-crimson-IRES-PuroR in pRRL.sin.cPPT.SFFV/E2-crimson-IRES-PuromycinR.WPRE with a BamHI/SalI fragment digested CD4-IRES-CXCR4 PCR fragment obtained from pMLV-CD4-IRES-CXCR4 (a gift from Prof. N. Sherer, Wisconsin University, USA). pRRL.sin.cPPT.SFFV/Firefly-IRES-PuromycinR.WPRE was obtained by amplification of Firefly by PCR from pGL4 (Promega) and cloned into BamHI-XhoI-digested pRRL.sin.cPPT.SFFV/E2-crimson-IRES-PuromycinR.WPRE. In some experiments, LVs without a selection marker were used: the IRES-PuromycinR cassette was removed by XhoI-SalI digestion and subsequent ligation, to obtain pRRL.sin.cPPT.SFFV/Firefly.WPRE and pRRL.sin.cPPT.SFFV/DDX42.WPRE. DDX42 K303E mutant was obtained by site-directed mutagenesis (by overlapping PCR using the aforementioned DDX42-forward and -reverse primers, respectively combined initially with reverse primer 5’-GGCTGCAGTTTCCCCACTACCTGTTTTGGCAATACC and forward primer 5’-GGTAGTGGGGAAACTGCAGCCTTCATTTGGCC). pRRL.sin.cPPT.SFFV/ACE2.WPRE has been described (48) (Addgene 145842). Flag-DDX42 and Flag-Firefly were amplified by PCR from the aforementioned LV plasmids and cloned into a NotI-XhoI-digested modified version of pCAGGS (49) to obtain pCAGGS/flag-DDX42.WPRE and pCAGGS/flag-Firefly.WPRE. The NL4-3/Nef–internal ribosome entry signal (IRES)-Renilla (NL4-3/Nef-IRES-Renilla) and the CCR5-version of this proviral clone were gifts from Prof. Sumit Chanda (20). Wild-type and Ba-L Env bearing HIV-1 NL4-3, IIIB and HIV-2 proviral clones have been described (50–52), as well as the transmitted founder HIV-1 molecular clones CH077.t, CH106.c, REJO.c (gifts from Prof. B. Hahn (27)) and HIV-2_ROD10_ and SIV_MAC239_ (53, 54). GFP-coding HIV-1 based LV system (i.e. p8.91 HIV-1 Gag-Pol, pMD.G, and GFP-coding minigenome), and HIV-2, FIV, and EIAV-derived, GFP coding LVs, as well as MLV-derived, GFP coding retroviral vectors have all been described (55, 56, 57, 58). The LINE-1 plasmid 99 RPS-GFP PUR (pRPS-GFP), 99 RPS-GFP JM111 PUR (pJM111) and pLRE3-GFP were developed by Prof. Kazazian’s lab (32, 59, 60).

### Cell lines

Human cell lines HEK293T, A549, U87-MG, TZM-bl were obtained from the ATCC and the AIDS reagent program, respectively. T98G cells were a gift from Prof. G. Kochs (Freiburg University, Germany), MDCK cells a gift from Prof. W. Barclay (Imperial College London, UK), Vero E6 cells (Merck) were a gift from Christine Chable (CEMIPAI, CNRS), respectively, Huh7.5.1 cells have been described (61), and the latter provided by Raphaël Gaudin. Human hepatocellular carcinoma Huh-7 cells (62) were kindly given by Annette Martin (Institut Pasteur, Paris). These cell lines were cultured in Dulbecco’s Modified Eagle Medium (DMEM) supplemented with 10% fetal bovine serum and 1 % penicillin-streptomycin (Thermofisher). T98G/Cas9 and U87-MG/Cas9 were obtained by transduction of T98G and U87-MG, respectively, with HIV-1-based LVs expressing the spCas9-P2A-Blasticidin cassette (pLentiCas9-Blast (17)). U87-MG/CD4/CXCR4 have been described (11) and were further modified to express Cas9 and Firefly using pLentiCas9-Blast and pRRL.sin.cPPT.SFFV/Firefly.WPRE, respectively. T98G/Cas9/CD4/CXCR4/Firefly were obtained by successive transductions of T98G/Cas9 with pRRL.sin.cPPT.SFFV/CD4-IRES-CXCR4.WPRE at a high MOI, and pRRL.sin.cPPT.SFFV/Firefly.WPRE, at a low MOI, respectively. Cell surface staining with anti-CD4 and CXCR4 antibodies (Miltenyi Biotec) confirmed than more than 95% cells were positive for both markers. A549 cells stably expressing ACE2 were generated by transduction with RRL.sin.cPPT.SFFV.ACE2.WPRE containing-vector.

For antibiotic selection, cells were treated with 10 μg/mL Blasticidin (InvivoGen), 1 mg/mL Zeocin (InvivoGen), 2 μg/mL Puromycin (Sigma-Aldrich), 250 μg/mL Hygromycin (Sigma-Aldrich), 1 mg/mL G418 (Sigma-Aldrich). When indicated, universal type 1 IFN (PBL Interferon Source) was added at 1000 U/mL for 16-24h prior to virus infection or RNA extraction, and AZT and 3TC (AIDS reagent program) at 10 μM for 2 h prior to infection.

### Primary cells

Blood from healthy donors was obtained from the Etablissement Français du Sang, under agreement n°21PLER2019-0106. Peripheral blood mononuclear cells (PBMCs) were isolated by centrifugation through a Ficoll® Paque Plus cushion (Sigma-Aldrich). Primary human CD4+ T cells and monocytes were purified by positive selection using CD3 and CD14 MicroBeads, respectively (Miltenyi Biotec), as previously described (11). Monocytes were incubated for 3 hours in serum-free Roswell Park Memorial Institute (RPMI) 1640 medium and further differentiated into macrophages by culture for 5-7 days in RPMI 1640 supplemented with 10% fetal calf serum, 1% penicillin-streptomycin and 100 ng/ml granulocyte-macrophage colony-stimulating factor (GM-CSF; Miltenyi). CD4+ T cells were cultured in RPMI supplemented with 10% fetal bovine serum and 1% penicillin-streptomycin, and stimulated for 48h with 10 μg/ml phytohemagglutinin (PHA) (Fisher Scientific) and 50 U/mL interleukin-2 (IL-2, Miltenyi Biotec) prior to electroporation.

### Genome-scale CRISPR/Cas9 screens

The plasmids coding GeCKO sub-libraries A and B were amplified and prepared according to the provided guidelines (Lentiviral Crispr Toolbox, Addgene). 60 million T98G/Cas9 cells were transduced with GeCKO LVs at a MOI of 0,1 to cover about 100-times the half-library complexity. After 48h, the cells were selected with puromycin, amplified for 12-15 days. 45 million cells were harvested and frozen down at −80°C for subsequent genomic DNA extraction, using the QIAamp DNA Blood Maxi Kit according to manufacturer’s instructions (Qiagen). In parallel, 60 million cells from the initial GeCKO populations were used for the screen. The cells were treated with 1000 U/mL IFN for 24h, infected with LVs coding a hygromycin resistance cassette. 48h later the cells selected with hygromycin and the surviving cells amplified. Two other rounds of IFN treatment, LV infection and antibiotic selection were subsequently performed with LVs coding a neomycin resistance cassette and a zeocin resistance cassette, respectively. The three time-selected populations were amplified and 45 million cells were harvested and stored at −80°C for subsequent genomic DNA extraction, as previously. After genomic DNA extraction, the sgRNA coding sequences integrated in the genomic DNA from the initial and 3-times selected populations were amplified by touch-down PCR and sequenced by Illumina deep sequencing. To this aim, 120 μg of genomic DNA was amplified using DNA Herculase II Fusion DNA polymerase (Agilent) in the presence of 2% DMSO, 1 mM of dNTPs; and 400 nM of the following primers: Forward-primer1: 5’-TCGTCGGCAGCGTCAGATGTGTATAAGAGACAGCTTGTGGAAAGGACGAAACACC-3’ for screen A or Forward-primer2: 5’-TCGTCGGCAGCGTCAGATGTGTATAAGAGACAGGATCTTGTGGAAAGGACGAAACACC-3’ used for screen B, together with reverse primer: 5’-GTCTCGTGGGCTCGGAGATGTGTATAAGAGACAAAGGTCCATTAGCTGCAAAGATTCCTCTC-3’). Briefly, after 5 minutes at 95°C, 14 cycles of pre-amplification were performed with a hybridization temperature decreasing by 0.5°C per cycle (30 sec at 95°C, 30 sec at 60°C, 30 sec at 72°C), followed by 30 cycles of amplification (30 sec at 95°C, 30 sec at 53°C, 30 sec at 72°C). 50 ng of each amplicon was dual indexed in a 5-cycle PCR reaction using the PCR module and indexed primers from the Nextera kit (Illumina). Resulting libraries were purified on AMPure XP magnetic beads (Beckman Coulter) using a 0,8X ratio and verified on Fragment Analyzer using the HS NGS fragment kit (Agilent). Libraries were quantified using microfluorimetry (QuBit, Invitrogen), mixed with a PhiX library (Illumina) and sequenced on one single read 50nt lane of Hiseq2500 using the rapid mode.

Image analyses and base calling were performed using the Illumina HiSeq Control Software and Real-Time Analysis component (v1.18.66.3). Demultiplexing was performed using Illumina’s conversion software (bcl2fastq 2.20). The quality of the raw data was assessed using FastQC (v0.11.5) from the Babraham Institute and the Illumina software SAV (Sequencing Analysis Viewer). Potential contaminants were investigated with the FastQ Screen(63) (v0.11.4) software from the Babraham Institute.

Sequencing reads were trimmed using Cutadapt (64) (v1.13), with options -g [primer sequence] -u [length of remaining 3’ bases] -e 0.2 -m 18 -l 20, to remove primer sequences and retrieve the 20 bases long sequences corresponding to sgRNAs. These retrieved sequences were then aligned to the GecKOv2 Human Library (A or B) reference sequences (keeping only non-duplicated sgRNA sequences, the duplicated ones being annotated) using Bowtie (65) (v1.2), with options -v 2 -norc -S. Resulting bam files were sorted and indexed using Samtools (66) (v1.5). Quantification of sgRNAs was done using Samtools idxstats. MAGeCK (18) (v0.5.7) was used to normalize (total count method) and identify enriched sgRNAs and genes in 3-times selected cell populations versus starting GeCKO transduced cells (mageck test command).

### Lentiviral and retroviral production

To produce lentiviral vector particles, HEK293T cells were transfected by polyethylenimine (PEI) co-transfection with miniviral, HIV-1 based genome coding plasmids (e.g. LentiCas9-Blast, LentiGuide-Puro or pRRL-SFFV), p8.91 (HIV-1 GagPol) and pMD.G (VSV-G) at a ratio of 1:1:0.5, respectively. The medium was replaced after 6h and viral particles were harvested 42h later, filtered, and directly used to transduce target cells (or stored at −80°C). After 4 to 6 hours, the transduction medium was replaced with complete DMEM, and the cells were treated 48h later with the relevant antibiotics. The HIV-2-, FIV-, EIAV-GFP coding LVs were produced using GFP-coding HIV-2-, FIV-, EIAV-based miniviral genomes, together with HIV-2-, FIV-, EIAV- GagPol, expression constructs and pMD.G at a ratio of 1:1:0.5. MIGR1 MLV-derived retroviral vectors were obtained with B-MLV Gag-Pol-expressing plasmid pCIG3B, the GFP-expressing minigenome pMIGR1 and pMD.G. at a ratio of 1:1:0.5, respectively and harvested as previously described.

HIV-1 Renilla and NL4-3 HIV-1 were produced by standard PEI transfection of HEK293T. When indicated, pMD.G was cotransfected with the provirus at a 3:1 ratio. The culture medium was changed 6h later, and virus-containing supernatants were harvested 42h later. Viral particles were filtered, purified by ultracentrifugation through a sucrose cushion (25% weight/volume in Tris-NaCl-EDTA buffer) for 75 min at 4°C and 28,000 rpm using a SW 32 TI rotor (Beckman Coulter), resuspended in serum-free RPMI 1640 or DMEM medium and stored in small aliquots at −80°C. Viral particles were titrated using an HIV-1 p24^Gag^ Alpha-Lisa kit and an Envision plate reader (Perkin Elmer) and/or by determining their infection titers on target cells.

### Lentiviral and retroviral infections

For infections with replication-competent HIV-1 Renilla or wild-type and/or VSV-G pseudotyped-HIV-1, target cells were plated at 2.5 × 10^4^ cells per well in 96-well plates or at 2 × 10^5^ cells per well in 12-well plates and infected for 24-48 h before lysis and Renilla (and Firefly) luciferase activity measure (Dual-Luciferase® Reporter Assay System Promega) or fixation with 2% paraformaldehyde (PFA)-PBS, permeabilization (Perm/Wash buffer, BDBiosciences) and intracellular staining with the anti-p24^Gag^ KC57-FITC antibody (Beckman Coulter), as described previously (67). For TZM-bl assays, the β-galactosidase activity was measured using the Galacto-Star™ system (ThermoFisher Scientific). For infections with lentiviral and retroviral vectors, target cells were plated at 2.5 × 10^4^ cells per well in 96-well plates the day prior to infection with vectors at the indicated MOIs, and the percentages of infected cells were scored by flow cytometry 24h later. For primary CD4+ T cell infections, 10^5^ cells were infected with 100 ng p24^Gag^ of HIV-1 Renilla for 24 h prior to lysis and luciferase activity measure. For MDM infections, 8 × 10^4^ cells were infected with 100 ng p24^Gag^ of a CCR5-tropic version of HIV-1 Renilla for 30 h prior to lysis and luciferase activity measure.

### Retrotransposon assays

For GFP-based retrotransposon assays, HEK293T cells (2 × 10^5^ cells) were co-transfected with either 1 μg of pJM111 (a negative control with two point mutations in ORF1 that abolish retrotransposition), pRPS-GFP or pLRE3-GFP with either 1 μg of pCAGGS-Flag-Firefly or pCAGGS-Flag-DDX42. At 7 days post-transfection, the percentage of GFP-expressing cells was scored by flow cytometry.

### CRISPR/Cas9 knock-out

For CRISPR/Cas9 knock-out in cell lines, Lentiguide-Puro LVs coding sgRNAs targeting the indicated genes or non-targeting sgRNAs were produced, and U87-MG Cas9/CD4/CXCR4/Firefly were transduced for 6 h before replacing the supernatants with fresh, complete medium. The transduced cells were selected with puromycin two days later and amplified for 12-15 days. For CRISPR/Cas9 knock-out in activated primary CD4+ T cells, 2 million cells per condition were washed with PBS1X, and electroporated using the 4d-Nucleofector® (Lonza) and the Amaxa P3 primary cell kit with 183 pmol of crispr/tracr RNA duplex (Alt-R CRISPR-Cas9 crRNA XT and tracrRNA XT, IDT®) and 61 pmol of Cas9 (Alt-R® S.p. Cas9 Nuclease V3, IDT®). After electroporation, the cells were incubated for 4 days at 37°C in X-VIVO15 medium (Lonza) supplemented with 1% pen/strep and IL-2 at 500 U/ml prior to cell counting and infection. The crRNA sequences of the sgDDX42-1, −2, and −3 were identical to the ones cloned in pLentiguide-Puro, and the crRNA of the sgDDX42-4 and sgDDX42-5 were pre-designed by IDT®, as follow: sg4-DDX42 5’-CGGAGATCTATTAACTGCTG-3’, sg5-DDX42 5’-GAGTTGGTGAGTTTTCAGC-3’.

### siRNA transfection

DDX42 and control knockdowns were achieved by transfecting the indicated siRNAs at 44nM, 14.2nM, and 100nM final in U87-MG and Huh-7 cells, TZM-bl cells and MDMs, respectively, with lipofectamine RNAimax (Thermofisher Scientific) according to the manufacturer’s instructions. The scramble siRNA controls used were universal siCTRL1 (SIC001) and siCTRL2 (SIC002) (Sigma-Aldrich) and the sequences of the siRNAs targeting DDX42 were siDDX42-1: 5’-CAGAAUGCCUGGUUUCGGA-3’ (SASI_Hs01_00119846, Sigma-Aldrich®), siDDX42-2: 5’-CUUACCUUGUGUUUGAUGA-3’ (SASI_Hs01_00119845, Sigma-Aldrich®), siDDX42-4: 5’-AUCUCGAAUACCCUUUACG-3’ (ID:136410, Ambion®).

### Cross-linking RNA immunoprecipitation

For LINE-1 RNA immunoprecipitation, HEK293T cells were co-transfected with equal amounts of pRPS-GFP and pCAGGS-Flag-DDX42 or - Flag-Firefly. For viral RNA immunoprecipitation, U87-MG cells were transduced with either Flag-Firefly or Flag-DDX42 coding lentiviral vectors (pRRL.sin.cPPT.SFFV/Flag-Firefly.WPRE and pRRL.sin.cPPT.SFFV/Flag-DDX42.WPRE, respectively) and infected with CHIKV at MOI 0.1, SARS-CoV-2 at MOI 0.13, or A/Victoria/3/75 IAV at MOI 0.1 for 24 h. 4 days post LINE-1-transfection or 24 h post-infection, cells were washed twice in PBS1X, incubated for 10 min with 0,1% formaldehyde in PBS1X, for 5 min in 250 mM Glycine and washed twice in cold PBS1X. Cells were lysed in RIPA buffer (50 mM Tris/HCl pH 7.5, 150 mM NaCl, 1% NP-40, 0.5% sodium deoxycholate, 1 mM EDTA, protease inhibitor cocktail and 40 U/mL RNasin). The lysates were clarified by centrifugation at 16,000*g*, for 10 min at 4 °C. Fractions of cell lysates were harvested at this stage to serve as controls for protein and RNA inputs (15%) and the rest was incubated with Flag-magnetic beads (ThermoFisher Scientific) for 2 h at 4 °C. The beads were washed 5 times in RIPA buffer and the immunoprecipitated proteins eluted using 150 μg/mL 3x Flag peptide (Sigma-Aldrich) in elution buffer (50 mM Tris/HCl pH 7.5, 75 mM NaCl, 1mM DTT, protease inhibitor cocktail and 40 U/mL RNasin) for 2 h. Fractions of eluates were harvested for immunoblot analysis (1/6^th^) and the rest subjected to RNA extraction (5/6^th^). RNA extractions were then performed using TRIzol (ThermoFisher Scientific).

### RNA quantification by RT-qPCR

To check silencing efficiency or measure gene induction after IFN treatment, 0,5-2 × 10^6^ cells were collected 2-3 days after siRNA transfection or 24h after IFN treatment or no treatment, and total RNAs were isolated using the RNeasy kit with on-column DNase treatment (Qiagen). cDNAs were generated using 250 ng RNA (High-Capacity cDNA Reverse Transcription Kit, Applied Biosystem, ThermoFisher Scientific, catalogue number 4368814) and analysed by quantitative (q)PCR using TaqMan gene expression assays (Applied Biosystem) specific for ACTB (Hs99999903_m1), GAPDH (Hs99999905_m1), and DDX42 (Hs00201296_m1). Triplicate reactions were run according to the manufacturer’s instructions using a ViiA 7 Real-Time PCR system. For relative quantification, samples were normalized to both ACTB and GAPDH mRNA expression and ΔΔCt analysis was performed.

For the measure of LINE-1 RNAs, 100 ng RNA (from cell extracts) or 25 μl of RNA extracted from the IP eluates (i.e. ~60% of the total amount of immunoprecipitated RNA) were reverse transcribed and analysed by qPCR using primers and probe specific for *ORF2*: ORF2-forward 5’-CACCAGTTAGAATGGCAATCATTAAA-3’, ORF2-reverse 5’-GGGATGGCTGGGTCAAATGG-3’ with ORF2-probe 5’-[FAM]-AGGAAACAACAGGTGCTGGAGAGGATGC-[TAMRA]-3. Absolute quantification was performed using a pRPS-GFP standard curve.

For the measure of SARS-CoV-2 replication, 3 × 10^5^ cells were harvested and total RNA was extracted using the RNeasy kit (Qiagen) employing on-column DNase treatment. 125 ng of cellular RNAs were used to generate cDNAs that were analysed by qPCR using RdRp primers and probe, as follow: RdRp_for 5’-GTGARATGGTCATGTGTGGCGG-3’, RdRp_rev 5’-CAAATGTTAAAAACACTATTAGCATA-3’, and RdRp_probe 5’-[FAM]-CAGGTGGAACCTCATCAGGAGATGC-[TAMRA]-3’ (68). pRdRp (which contains an RdRp fragment amplified from SARS-CoV-2-infected cell RNAs (48)) was diluted in 20 ng/ml salmon sperm DNA to generate a standard curve to calculate relative cDNA copy numbers and confirm the assay linearity (detection limit: 10 molecules of RdRp per reaction).

For the measure of the amounts of viral RNAs in the RNA immunoprecipitation experiments, 100 ng RNA from cell lysates (input) or 25 μl of RNA extracted from the IP eluates (i.e. ~60% of the total amount of immunoprecipitated) were reverse transcribed using the High-Capacity cDNA Reverse Transcription Kit as above. For SARS-CoV-2, the cDNAs were analysed by RdRp RT-qPCR. For CHIKV RNA, the following primers and probe were used for the qPCR reactions: E1-C21-forward 5’-ACGCAGTTGAGCGAAGCAC-3’, E1-C21-reverse 5’-CTGAAGACATTGGCCCCAC-3’ (69), and E1-C21-probe 5’-[FAM]-CTCATACCGCATCTGCATCAGCTAAGCTCC-[TAMRA]-3’. pE1 (which contains an E1 fragment amplified from CHIKV-infected cell RNAs using primers E1-C21 forward and E1-C21 reverse and cloned into pPCR-Blunt II-TOPO) was used to generate a standard curve and ensure the linearity of the assay (detection limit: at least 10 molecules per reaction). For A/Victoria/3/75 IAV RNA, the following primers and probe, specific for the PA segment, were used, as follow: PA-forward 5’-TTGCTGCACAGGATGCATTA-3’, PA-reverse 5’-AGATTGGAGAAGACGTGGCT-3’ and PA-probe 5’-[FAM]-TGGCTCTGCAATGGGACACCTCTGC-[TAMRA]-3’. pPolI-RT-Victoria-PA (70) was used to generate a standard curve and ensure the linearity of the assay (detection limit: at least 10 molecules per reaction).

### Quantification of HIV-1 DNAs

To measure HIV-1 cDNAs, 2 × 10^5^ cells transfected with a control siRNA or siRNAs targeting DDX42 were plated in 24-well plates, and treated or not with 10 μM AZT and 3TC 1-2 h prior to infection. The cells were infected with NL4-3 HIV-1 (60 ng p24^Gag^) for 2 h at 37°C, washed with PBS1X and incubated in complete DMEM before being harvested at the indicated times. Cell pellets were frozen at −80°C after two washes in PBS1X. Total DNA extraction was performed using the DNeasy kit (Qiagen) according to the manufacturer’s instructions, and a DpnI-treatment step was performed prior to qPCR. Strong stop reverse transcription products were detected using forward primer oHC64 5’-TAACTAGGGAACCCACTGC-3’ and reverse primer oHC65 5’-GCTAGAGATTTTCCACACTG-3’, 2^nd^ strand transfer product using oHC64 and oSA63R 5’-CTGCGTCGAGAGATCTCCTCTGGCT-3’, together with oHC66 probe 5’-[FAM]-ACACAACAGACGGGCACACACTA-[TAMRA]-3’. 2-LTR circular forms were detected using 2LTR-forward 5’-GTAACTAGAGATCCCTCAG-3’ and 2LTR-reverse 5’-TGGCCCTGGTGTGTAGTTC-3’ together with 2LTR-probe 5′-[FAM]-CTACCACACACAAGGCTACTTCCCTGAT-[TAMRA]-3’. Integrated viral DNA was analysed using an Alu qPCR as described before (11). Briefly, a preamplification of 16 cycles was performed (15 sec at 94°C, 15 sec at 55°C, 100 sec at 68°C) with Platinum Taq DNA High Fidelity polymerase (Invitrogen) using 100 nM of genomic Alu forward primer 5′-GCCTCCCAAAGTGCTGGGATTACAG and 600 nM of U3-reverse primer 5’-CTTCTACCTTATCTGGCTCAAC-3’. The pre-amplification step was performed on serial dilutions of all the DNA samples, as well as of a positive control (total DNA from U87-MG infected with a high input of NL4-3), to ensure the linearity of the assay. Background levels were assessed using linear, one-way amplification by performing the pre-amplification PCR with the U3-reverse primer alone. Then a qPCR was performed on pre-amplification products using U3-forward primer 5’-TCTACCACACACAAGGCTAC-3’ and U3-reverse primer with the U3 probe 5’-[FAM]-CAGAACTACACACCAGGGCCAGGGGTCA-[TAMRA]-3’. qPCR reactions were performed in triplicates, in Universal PCR master mix using 900nM each primer and 250nM probe with the following program: 10 min at 95°C followed by 40 cycles (15 sec at 95°C and 1 min at 60°C). pNL4-3 or pTOPO-2LTR (generated by pTOPO cloning of a 2-LTR circle junction amplified from NL4-3 infected cells, using oHC64 and U3-reverse primers into pCR™2.1-TOPO™) were diluted in 20 ng/ml of salmon sperm DNA to create dilution standards used to quantify relative cDNA copy numbers and confirm the linearity of all assays.

### Proximity Ligation assays (PLAs)

The proximity ligation assays were performed using the Duolink® in situ Detection Reagents (Sigma-Aldrich, DUO92014). For PLA with HIV-1, MDMs were plated in 24-well plates with coverslips pre-treated with poly-L-lysin (Sigma-Aldrich) and infected with 1 μg p24^Gag^ of HIV-1 NL4-3 (Ba-L Env) or mock-infected. For PLA with SARS-CoV-2, A549-ACE2 cells were plated in 24-well plates with coverslips and infected at an MOI of 0,1. 24 h later, the cells were fixed with 2-4% paraformaldehyde in PBS1X for 10 min, washed in PBS1X and permeabilized with 0.2% Triton X-100 for 10 min. After a couple of washes in PBS1X, either NGB buffer (50 mM NH_4_Cl, 2% goat serum and 2% bovine serum albumin in PBS) or Duolink® blocking solution was added for 1h. Cells were incubated with mouse AG3.0 anti-HIV-1 Capsid antibody (National Institutes of Health (NIH) AIDS Reagent Program #4121), or J2 anti-dsRNA antibody (SCICONS), or anti-SARS-CoV-2 Nucleoprotein (N; BioVision), and rabbit anti-DDX42 antibody (HPA023571, Sigma-Aldrich) diluted in NGB buffer or in Duolink® blocking solution for 1h. After 2 washes in PBS1X, the cells were incubated with the DUOLINK® in situ PLA® Probe Anti-rabbit minus (DUO92006) and DUOLINK® in situ PLA® Probe Anti-mouse plus (DUO92001) for 1h at 37°C. After 2 washes in PBS1X, the ligation mix was added for 30 min at 37°C. After 2 washes in PBS1X, the cells were incubated with the amplification mix for 100 min at 37°C followed by 3 washes in PBS1X. In the case of SARS-CoV-2 infection, an additional staining was performed by incubating cells in an anti-mouse Alexa Fluor secondary antibody. Finally, the cells were washed twice with PBS1X and stained with Hoechst at 1 μg/mL for 5 min, washed again and the coverslips mounted on slides in Prolong mounting media (ThermoFisher Scientific). Z-stack images were acquired using an LSM 880 confocal microscope (ZEISS) using a 63x lens. PLA punctae quantification was performed using the FIJI software (71). Briefly, maximum z-projections were performed on each z-stack and the number of nuclei per field were quantified. Then, by using a median filter and thresholding, PLA punctae were isolated and quantified automatically using the Analyse Particles function. To obtain a mean number of dots per cell, the number of PLA dots per field were averaged by the number of nuclei. In the case of SARS-CoV-2 infection, the infected cells were identified using N or dsRNA immunofluorescence staining. For representative images, single cells were imaged using a LSM880 confocal microscope coupled with an Airyscan module. Processing of the raw Airyscan images was performed on the ZEN Black software.

### Immunoblot analysis

Cell pellets were lysed in sample buffer (200 mM Tris-HCl, pH 6.8, 5.2% SDS, 20% glycerol, 0.1% bromophenol blue, 5% β-mercaptoethanol), resolved by SDS–PAGE and analysed by immunoblotting using primary antibodies specific for human DDX42 (HPA023571, Sigma-Aldrich), Flag (M2, Sigma-Aldrich) and Actin (A1978, Sigma-Aldrich), followed by secondary horseradish peroxidase-conjugated anti-mouse or anti-rabbit immunoglobulin antibodies and chemiluminescence (Bio-Rad). Images were acquired on a ChemiDoc™ gel imaging system (Bio-Rad).

### IAV production and infection

We have described previously IAV NanoLuciferase reporter virus generation (47). Stocks were titrated by plaque assays on MDCK cells. IAV challenges were performed in serum-free DMEM for 1 h and the medium was subsequently replaced with DMEM containing 10%. IAV infection experiments were performed in triplicates in 96-well plates with cultures maintained for 16 h post-challenge. NanoLuciferase activity was measured with the Nano-Glo assay system (Promega), and luminescence was detected using a plate reader (Infinite® 200 PRO, Tecan).

### VSV production and infection

A VSV-G pseudotyped-VSV-Δenv reporter virus, coding both GFP and Firefly Luciferase, was obtained from Gert Zimmer. The virus was amplified on pMD.G transfected HEK293T and titrated thanks to the GFP reporter gene. For infection, 2.5 × 10^4^ cells per well in 96-well plates were infected at the indicated MOIs. 24h after infection, cells were lysed and Firefly luciferase activity was measured (Firefly luciferase Assay System Promega).

### Measles virus production and infection

Measles virus GFP strain (MeV-GFP), which was kindly provided by F. Tangy (Institut Pasteur, Paris), was previously described (73). Viral stocks were produced on Vero NK cells. After 4 days of infection, supernatant was collected and then centrifugated to eliminate dead cells or fragments. Stocks were tittered using median tissue culture infectious dose assays (TCID_50_) on Vero NK cells. Cells were infected with 10-fold serial dilutions of viral stocks and incubated for 7 days. Cells were then washed with PSB and fixed with 3% formaldehyde crystal violet during 30 min and finally rinsed with water. For infections, Huh-7 cells were infected at the indicated multiplicity of infection (MOI) in DMEM without FBS for 2 h in small volume of medium to enhance contacts with the inoculum and the cells. After 2 h, the viral inoculum was replaced with fresh DMEM 10% FBS 1% P/S. 24 h post-infection the cells were harvested and samples separated in half for Western blot and flow cytometry analysis.

### ZIKV production and infection

The nanoluciferase expressing ZIKV construct has been described (74). The corresponding linearized plasmid was transcribed in vitro using the mMESSAGE mMACHINE™ SP6 transcription kit (Thermofischer Scientific) and HEK293T cells were transfected with the transcribed RNA. After 7 days, supernatants were harvested, filtered and stock titers were determined by plaque assays on Vero cells. For infections, 2.5 × 10^4^ cells per well in 96-well plates were infected, at the indicated MOIs. 24h after infection, cells were lysed and Nanoluciferase activity was measured using the Kit Nano Glo luciferase Assay (Promega).

### CHIKV production and infection

The Nanoluciferase luciferase coding CHIKV construct was a gift from Andres Merits. The linearized plasmid coding CHIKV genome was transcribed with the mMESSAGE mMACHINE SP6 kit (Thermofischer Scientific) and 5 × 10^5^ HEK293T were transfected with 1-4 μg of transcribed RNA, using Lipofectamine 2000 (Thermofischer Scientific). After 24h, supernatants were harvested, filtered and viruses were then amplified on baby hamster kidney (BHK21) cells. Stock titers were determined by plaque assays on Vero cells. For infections, 2.5 × 10^4^ cells per well in 96-well plates were infected at the indicated MOIs. 24h after infection, cells were lysed and Nanoluciferase activity was measured.

### SARS-CoV-2 production and infection

The SARS-CoV-2 BetaCoV/France/IDF0372/2020 isolate was supplied by Pr. Sylvie van der Werf and the National Reference Centre for Respiratory Viruses hosted by Institut Pasteur (Paris, France). The patient sample from which strain BetaCoV/France/IDF0372/2020 was isolated was provided by Dr. X. Lescure and Pr. Y. Yazdanpanah from the Bichat Hospital, Paris, France. The virus was amplified in Vero E6 cells (MOI 0,005) in serum-free media supplemented with 0,1 μg/ml L-1-p-Tosylamino-2-phenylethyl chloromethylketone (TPCK)-treated trypsin (Sigma-Aldrich). The supernatant was harvested at 72 h post infection when cytopathic effects were observed (with around 50% cell death), cell debris were removed by centrifugation, and aliquots stored at −80C. Viral supernatants were titrated by plaque assays in Vero E6 cells. Typical titers were 5.10^6^ plaque forming units (PFU)/ml. Infections of A549-ACE2 cells were performed at the indicated multiplicity of infection (MOI; as calculated from titers obtained in Vero E6 cells) in serum-free DMEM and 5% serum-containing DMEM, respectively. The viral input was left for the duration of the experiment and cells lysed at 48 h post-infection for RT-qPCR analysis.

### Statistical analyses

Statistical analyses were performed with GraphPad Prism. Analysis types are mentioned in Fig. legends and all comparisons are relative to the indicated controls. For data with experimental factors greater than two, multiple linear regression was performed. For data with two categorical factors, ANOVA was used, and repeated measures ANOVA when a pairing factor was present. Simple linear regression was used when the relationship between a continuous factor and a continuous response variable was investigated. P values are denoted as follow: ns not significant, p<0.05 *, p<0.01 **, p<0.001 ***, p<0.0001****.

## Data availability

The datasets generated during and/or analysed during the current study are available from the corresponding author on reasonable request.

## Requests for materials

Requests for material should be addressed to Caroline Goujon at the corresponding address above.

## Acknowledgements

We wish to thank Tom Doyle and Chad Swanson for their useful comments on the manuscript, and Matthieu Lewis, Nadine Laguette, Nathalie Arhel, Juliette Fernandez, Jean-Luc Battini, Georg Kochs, Andres Merits, Frédéric Tangy, Sylvie van de Werf and Nathan Sherer for the generous provision of reagents, protocols and/or for helpful discussions. We are grateful to Nicolas Manel and Helena Izquierdo-Fernandez for sharing their protocol for efficient knockdown in primary CD4+ T cell using CRISPR/Cas9 RNP electroporation. We are grateful to Myriam Boyer, Stéphanie Viala, Baptiste Monterroso and Orestis Faklaris from the imaging and flow cytometry facility MRI, for advice with flow cytometry and confocal microscopy, respectively. This work was supported by the Institut National de la Santé et de la Recherche Médicale (INSERM) (to CG), the ATIP-Avenir programme (to CG), institutional funds from the Centre National de la Recherche Scientifique (CNRS) and Montpellier University, France REcherche Nord&Sud Sida/HIV et Hépatites (ANRS) (ECTZ21792 to CG; ECTZ35478 to OM), Sidaction (to CG), a Starting Grant from the European Research Council (759226, ANTIViR) (to CG), PhD studentships from the Ministry of Higher Education and Research (to BB, JM and AR), and a 4^th^ year PhD studentship from the Fondation pour la Recherche Médicale (to BB, FDT201904008024). SR and HP acknowledge financial support from the France Génomique National infrastructure, funded as part of “Investissement d’Avenir” program managed by Agence Nationale de la Recherche (contract ANR-10-INBS-09). We acknowledge the imaging facility MRI, member of the national infrastructure France-BioImaging supported by the French National Research Agency (ANR-10-INBS-04) and the BSL-3 facility CEMIPAI (UAR 3725 CNRS Montpellier University).

## Author contributions

B.B. and CG designed the study, analysed the data and wrote the manuscript. B.B. and C.G. performed the whole-genome screens and candidate validation. B.B. carried out most of the experiments, with technical assistance from A.R., F.G.d.G, J.M., M.T., W.D., A.M., A.L.C.V., M.A.A., and O.M.; V.C. provided some of the lentiviral vector stocks; E.B. and L.B. performed CHIKV infections, N.G. performed ZIKV-Nluc infections, S.G. and N.J. performed MeV, ZIKV PF13, DENV-2 and YFV infections; M.L and E.R analysed the RNA-seq data; R.S. performed the statistical analyses; H.P. and S.R. performed the Illumina sequencing and MaGECK analyses, respectively. All authors have read and approved the manuscript.

## Conflicts of interest statement

The authors have no conflicts of interest to declare in relation to this manuscript.

## Notes

### Competing Interest Statement

The authors have declared no competing interest.

### Summary of Updates

New data in this version of our manuscript strongly support the hypothesis of a direct mode of action of DDX42 on retroelements and viruses. Indeed, RIP experiments show a specific interaction between DDX42 and RNAs from 2 sensitive viruses, SARS-CoV-2 and CHIKV, but not with RNA from an insensitive virus (IAV). RNA-seq analyses show that DDX42 depletion does not have a generalized impact on the transcriptome that could have explained the observed antiviral phenotypes. Moreover, we have expanded the analysis of DDX42 antiviral breadth, by adding data on negative-strand RNA virus VSV and on 3 additional positive-strand RNA viruses, the flaviruses YFV and DENV-2 and the seasonal coronavirus HCoV-229E.

