## Supplementary Figures and Methods for "The DEAD box RNA helicase DDX42 is an intrinsic inhibitor of positive-strand RNA viruses"

### SUPPLEMENTARY INFORMATION

#### Supplementary Figures

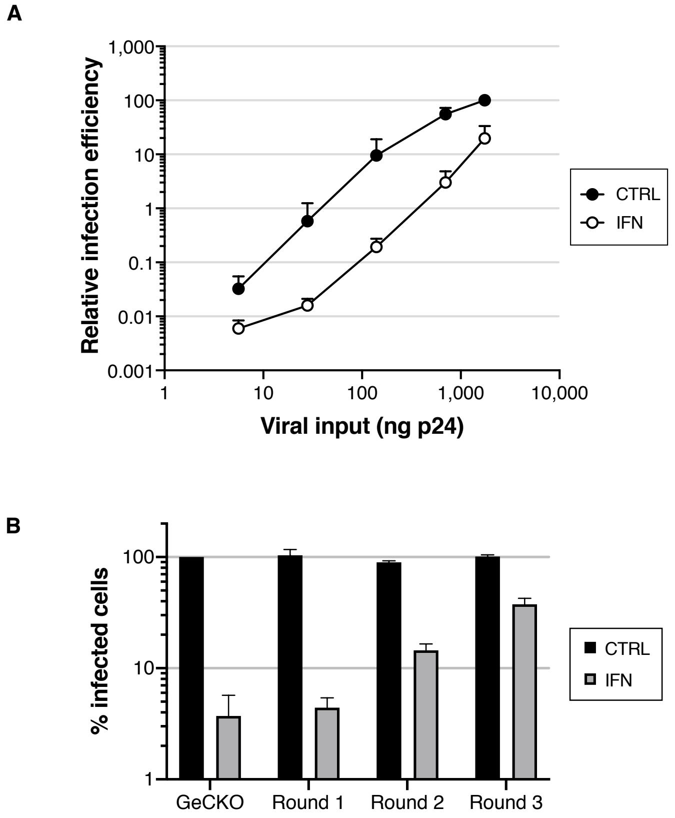


**Figure S1. Related to Figure 1.**

**A. IFN pre-treatment potently inhibits HIV-1 infection in T98G cells and is, at least partially, saturable.** T98G/Cas9/CD4/CXCR4/Firefly cells were pre-treated with IFN for 24 h prior to infection with increasing doses of NL4-3 Renilla (indicated in ng $p24^{Gag}$). Renilla activity was normalized to Firefly activity and the relative infection efficiencies are shown. Data represent the average and standard deviation (s.d.) of 3 independent experiments.

**B. GeCKO screen validation.** GeCKO control cells and enriched cells from 3 successive rounds of selection (Round 1, 2, and 3, as indicated) were treated with IFN or not and infected with GFP-expressing lentiviral vectors. The percentage of infected cells was evaluated by flow cytometry 2 days post-infection. Data represent the average and s.d. of 2 independent experiments.

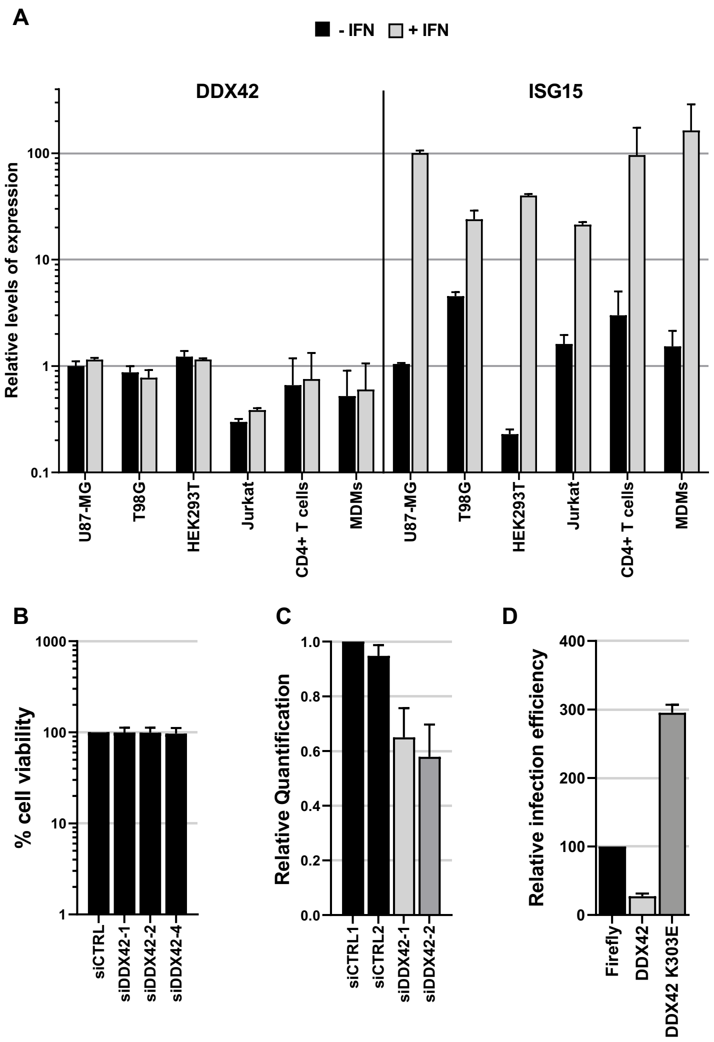


**Figure S2. Related to Figure 2.**

**A. DDX42 is not an ISG**. The indicated primary cells or immortalized cell cultures were treated with IFN for 24 h or left untreated. RNA was subsequently extracted and DDX42 and ISG15 (a prototype ISG) mRNA levels were quantified by RT-qPCR; Actin B and GAPDH were used as endogenous controls. The bar chart shows the relative levels of expression of DDX42 and ISG15 in the presence or absence of IFN treatment. Data represent the mean ± S.E.M of 3 independent experiments.

**B. Viability assay in siRNA transfected cells.** Cell viability of siRNA-transfected U87-MG/CD4/CXCR4 cells was assessed 72h post-transfection by measuring ATP levels. Data represent the mean ± S.E.M of 3 independent experiments.

**C. DDX42 silencing efficiency in monocyte-derived macrophages.** siRNA-transfected MDMs were harvested 48h post-transfection for RNA extraction and quantification of DDX42 mRNA levels by RT-qPCR. Actin and GAPDH were used as endogenous controls. Data represent the mean ± S.E.M of 3 independent experiments performed with cells from 3 different blood donors (parallel samples from Figure 2E).

**D. Ectopic expression of DDX42 K303E mutant increases HIV-1 infection.** U87-MG/CD4/CXCR4 cells were transduced with lentiviral vectors expressing either Firefly (negative control), WT DDX42 (DDX42) or a motif I point mutant, which has an impaired ATPase activity (DDX42 K303E). Transduced cells were infected with NL4-3 Renilla and the infection efficiency was assessed 24h later by measuring Renilla activity. Data represent the mean ± S.E.M of 4 independent experiments.

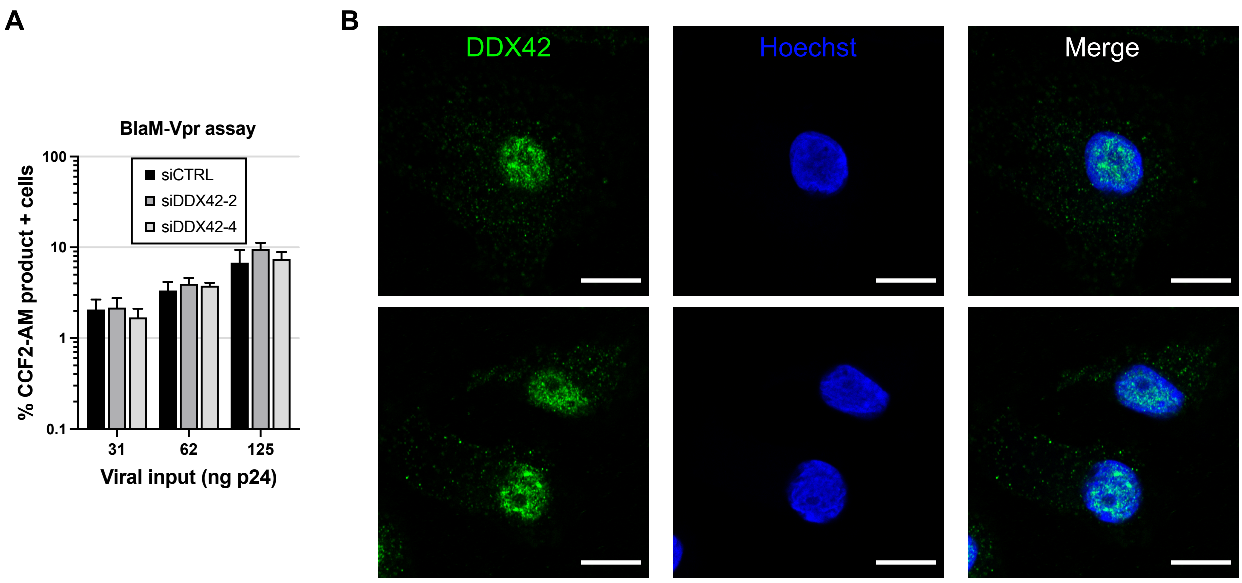


**Figure S3. Related to Figure 3.**

**A. DDX42 depletion does not affect HIV-1 entry.** DDX42-depleted U87-MG/CD4/CXCR4 cells were infected with the indicated amounts of viruses carrying the β-lactamase (BlaM)-Vpr fusion protein for 3 h. The cells were subsequently loaded with CCF2-AM substrate dye for 2h, washed extensively and incubated for another 16 h for the reaction to develop. Cells positive for the CCF2-AM product were scored by flow cytometry. Data represent the mean ± S.E.M of 3 independent experiments.

**B. DDX42 localization in MDMs.** MDMs were fixed, endogenous DDX42 and the nuclei were visualized using DDX42-specific antibodies and Hoechst staining, respectively, and confocal microscopy. Representative images are shown. Scale bar, 10 μm.

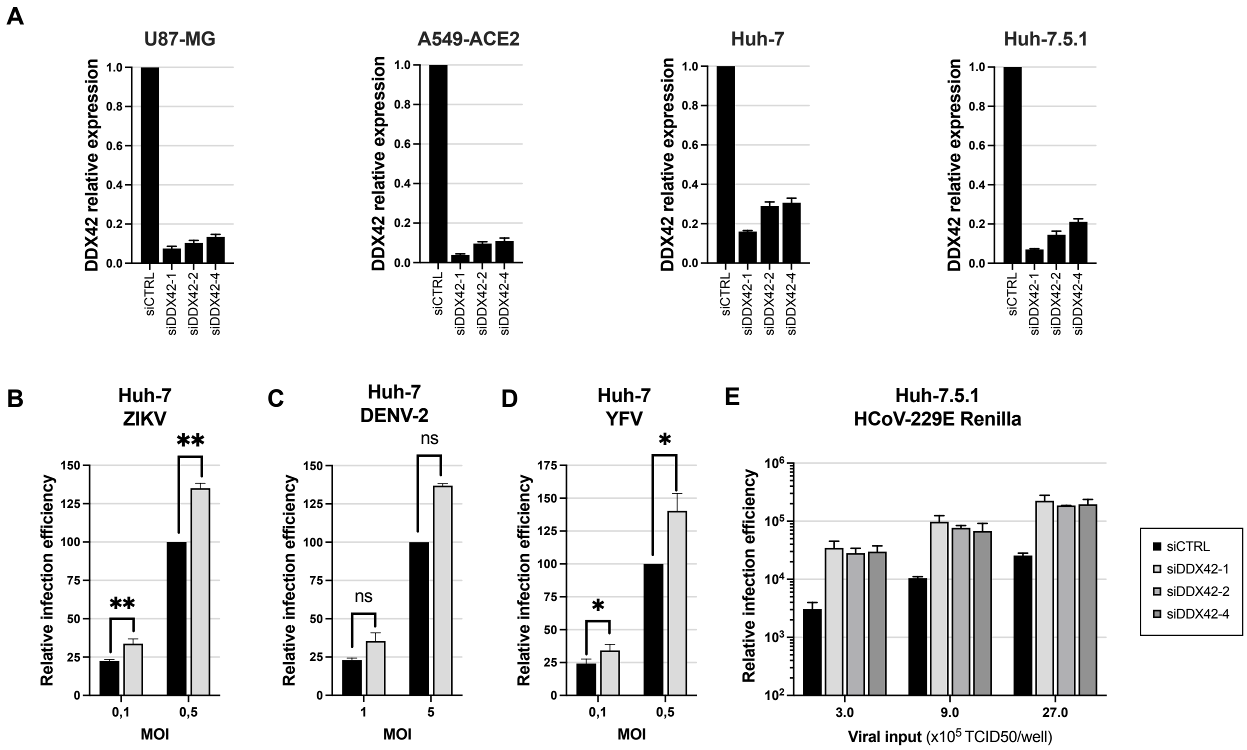


**Figure S4. Related to Figure 4.**

**A.** DDX42 silencing efficiency in U87-MG, A549-ACE2, Huh-7 and Huh-7.5.1 cells.

**B.** Relative ZIKV PF13 infection efficiency in siRNA-transfected Huh-7 cells analysed by flow cytometry.

**C.** Relative DENV-2 infection efficiency in siRNA-transfected Huh-7 cells analysed by flow cytometry.

**D.** Relative YFV infection efficiency in siRNA-transfected Huh-7 cells analysed by flow cytometry.

**E.** Relative HCoV-229E Renilla infection efficiency in siRNA-transfected Huh7.5.1 cells analysed by Renilla signal monitoring.

**A-E**. Mean ± SEM of 3 independent experiments (4 for silencing efficiency in A549-ACE2 cells). Two-way ANOVA with Sidak's test on log10 transformed data.

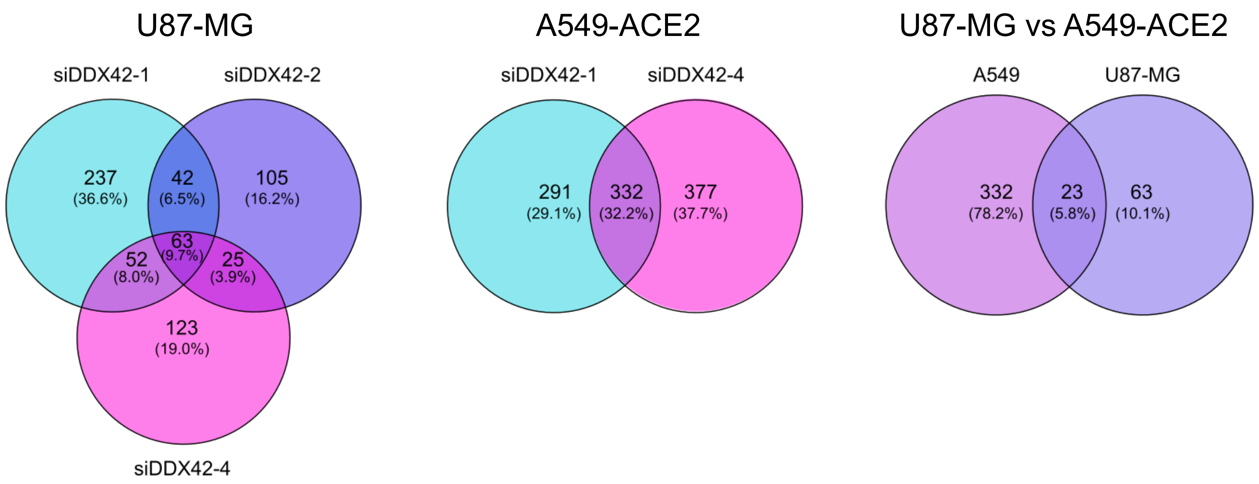


**Figure S5.**

Venn diagrams showing the Differentially Expressed Genes (DEGs) overlap between siRNA conditions in U87-MG cells, A549 cells and DEGs overlapping between U87-MG and A549 cells using cutoff criteria of log2 fold change (log2FC) >1 and p value <0.05.

See **Supplemental File 1** for the identity of the DEGs.

**Supplemental methods**

Only the methods specific for the supplemental Figures are described here.

**Plasmids**. pBlaM-Vpr and pAdVAntage have been described (1).

**Lentiviral production**. β-lactamase-Vpr (BlaM-Vpr)-carrying viruses, bearing the wild-type Env, were produced by co-transfection of HEK293T cells with the NL4-3/Nef-IRES-Renilla provirus expression vector, pBlaM-Vpr and pAdVAntage at a ratio of 4:1:0.5, as previously described (1). Viral particles were titrated using an HIV-1 $p24^{Gag}$ Alpha-Lisa kit and an Envision plate reader (Perkin Elmer).

**BlaM-Vpr assay for HIV-1 entry.** This assay was performed as described previously (2). Briefly, 2 x $10^5$ U87-MG/CD4/CXCR4 cells were plated in 24-well plates and incubated with BlaM-Vpr carrying NL4-3 particles (31, 62, 125 ng $p24^{Gag}$) or mock-infected for 3 h at 37°C. The cells were then washed once in $CO_2$-independent medium and loaded with CCF2-AM substrate-containing solution (ThermoFisher Scientific) for 1 h at room temperature before 2 washes and incubation at room temperature for 16 h in development medium ($CO_2$-independent medium containing 2,5 mM probenecid). Finally, the cells were trypsinized, washed and fixed in 1% paraformaldehyde (PFA)-PBS1X before analysis with a FACSCanto™ II (Becton Dickinson).

**Viruses and infection.** The Yellow Fever Virus (YFV) Asibi strain was provided by the Biological resource Center of Institut Pasteur. Stocks were produced on Vero NK cells. After 3 days of infection, viruses were concentrated by polyethylene glycol 6000 (PEG) precipitation and purified by centrifugation in a discontinuous gradient of sucrose. The Dengue 2 strain Malaysia SB8553 (DENV-2) was obtained from the Centro de Ingeniería Genética y Biotecnología (CIGB), Cuba. Stocks were generated on Vero NK cells. After 4 days of infection, viruses were concentrated by PEG 6000 precipitation. The Zika strain PF13 (kindly provided by V. M. Cao-Lormeau and D. Musso, Institut Louis Malardé, Tahiti Island, French Polynesia) was isolated from a viremic patient in French Polynesia in 2013. Stocks were produced on C6-36 cells. After 2 days of infection, viruses were concentrated by PEG 6000 precipitation and purified by centrifugation in a discontinuous gradient of sucrose. YFV Asibi and ZIKV titers were assessed by plaque assays using Vero NK cells, as described previously (3). DENV-2 was tittered by in cell western assays on Vero cells. Cells were fixed with PFA 4% during 30 min at room temperature (RT), then washed in PBS and permeabilized with 0,5% triton in PBS (Sigma-Aldrich) during 10 min at RT. Cells were then incubated 0,1% Tween in PBS (Sigma-Aldrich) containing 5% BSA (Sigma-Aldrich) during 1 h at RT prior to incubation with mouse anti-Env 4G2 antibodies overnight at 4°C. After 1h of incubation with the secondary antibodies, cells were revealed with an Odyssey CLx infrared imaging system (Li-Cor Bioscience). Cells were infected at the indicated multiplicity of infection (MOI) in DMEM without FBS for 2 h in small volume of medium to enhance contacts with the inoculum and the cells. After 2 h, the viral inoculum was replaced with fresh DMEM 10% FBS 1% P/S. 24 hours post-infection the cells were harvested and samples separated in half for Western blot and flow cytometry analysis. For the latter, cells were fixed and permeabilized using BD Cytofix/Cytoperm (Fisher scientific) for 30 min on ice (all the following steps were performed on ice and centrifuged at 4°C) and then washed tree times with wash buffer. Cells infected with YFV, ZIKV and DENV-2 were incubated with the pan-flavivirus anti-Env 4G2 antibody for 1 h at 4°C and then with Alexa 488 anti-mouse IgG secondary antibodies (Thermo Fisher) for 45 min at 4°C in the dark. Data were acquired with an Attune NxT Acoustic Focusing Cytometer (Life technologies) and analyzed using FlowJo software.

HCoV-229E-Renilla was a gift from Volker Thiel (4) and was amplified for 5-7 days at 33°C in Huh7.5.1 cells (5), in 5% FCS-containing DMEM. Virus stock concentration was assessed in Huh7.5.1 cells and titer was determined at $10^9$ TCID50/mL. Cell were infected at the indicated TCID50s in DMEM 5% FCS at 33°C. 24 h later, cells were lysed and Renilla activity was measured using the Renilla Luciferase Assay System kit (Promega).

**Cell viability assays**. Cell viability was assessed in siRNA-transfected cells by measuring cellular ATP levels 72 h post-transfection using the CellTiter-Glo® Luminescent Cell Viability Assay kit (Promega) according to manufacturer's instructions.

**Quantification of mRNA expression**. For all figures, see the main material and methods; For Fig. S2: a TaqMan gene expression assays (Applied Biosystem) specific for ISG15 was used: Hs00192713_m1.

For the RNA extraction and RT-qPCR analysis specifically in Huh-7 cells, total RNAs were extracted from cell lysates using the NucleoSpin RNA II kit (Macherey-Nagel), following the manufacturer's protocol. Equal amounts of purified total RNA were used to synthetized first strand cDNA using random hexamers (Thermo Fisher) and the ReverAid H Minus Moloney murine leukemia virus (M-MuLV) reverse transcriptase (Thermo Fisher). Quantitative real-time PCR was performed on a real-time PCR system (Quant Studio 6 Flex from Applied Biosytems) with SYBR green PCR master mix (Life Technologies). Data were analyzed with the $\Delta\Delta$ CT method. All the samples were performed in technical triplicate and normalized to GAPDH (glyceraldehyde-3-phosphate dehydrogenase) as endogenous reference control. The primer sequences used for Fig. S4 were as follow: GAPDH Forward: 5'-GGTCGGAGTCAACGGATTTG-3', reverse: 5'-ACTCCACGACGTACTCAGCG-3'; DDX42 Forward: 5'-GGCCTATACCCTACTCACTCCC-3', reverse: 5'-CCACCAATGTTCAGCTTTTTTCC-3'.

**Preparation of RNA-seq libraries.** siRNA transfected U87-MG/CD4/CXCR4 and A549-ACE2 RNA extracts from three independent experiments were used for RNA-seq library preparation. After determining sample RNA integrity numbers using a 2100 Bioanalyzer (Agilent), ribosomal RNAs were depleted using the QIAseq FastSelect -rRNA HMR Kit (Qiagen) and libraries were prepared according to manufacturer's instructions using the QIAseq Stranded Total RNA Lib Kit (Qiagen). Libraries were quantified using a TapeStation D1000 ScreenTape. Equimolar amounts of each library were then mixed and sequenced on 2 lanes (2x150 bp) on the Illumina HiSeq 3000/4000 platform (GENEWIZ).

**Analysis of high-throughput sequencing reads.** Sequenced reads were filtered by quality and sequence adaptors removed using fastp v0.20.1 (https://github.com/OpenGene/fastp)(6) with following parameters “fastp --qualified_quality_phred 20 --disable_length_filtering --detect_adapter_for_pe”. Reads were pseudo-mapped against human cDNA sequenced downloaded from Gencode database (https://www.gencodegenes.org/) and transcripts abundance estimated with Kallisto v0.46.2 (https://pachterlab.github.io/kallisto/about)(7) with parameters "--bias --bootstrap-samples 100".

**Differential analysis with DESeq2.** Differential expressed genes upon siRNA transfection were obtained using DESeq2 (8) (version 1.32.0) in R (version 4.1.0). Briefly, transcript estimations were transformed in gene counts with tximport package (9) and the differential expression obtained with the model design “~ condition + replicate”.

**Supplementary Information References**

6. Chen,S., Zhou,Y., Chen,Y. and Gu,J. (2018) fastp: an ultra-fast all-in-one FASTQ preprocessor. *Bioinformatics*, 34, i884–i890.

7. Bray,N.L., Pimentel,H., Melsted,P. and Pachter,L. (2016) Near-optimal probabilistic RNA-seq quantification. *Nat. Biotechnol.*, 34, 525–527.

8. Love,M.I., Huber,W. and Anders,S. (2014) Moderated estimation of fold change and dispersion for RNA-seq data with DESeq2. *Genome Biol.*, 15, 550.

9. Soneson,C., Love,M.I. and Robinson,M.D. (2015) Differential analyses for RNA-seq: transcript-level estimates improve gene-level inferences. *F1000Research*, 4, 1521.
